# Single-cell analysis of Foxp1-driven mechanisms essential for striatal development

**DOI:** 10.1101/611780

**Authors:** Ashley G. Anderson, Ashwinikumar Kulkarni, Matthew Harper, Genevieve Konopka

## Abstract

The striatum is a critical forebrain structure for integrating cognitive, sensory, and motor information from diverse brain regions into meaningful behavioral output. However, the transcriptional mechanisms that underlie striatal development and organization at single-cell resolution remain unknown. Here, we show that Foxp1, a transcription factor strongly linked to autism and intellectual disability, regulates organizational features of striatal circuitry in a cell-type-dependent fashion. Using single-cell RNA-sequencing, we examine the cellular diversity of the early postnatal striatum and find that cell-type-specific deletion of *Foxp1* in striatal projection neurons alters the cellular composition and neurochemical architecture of the striatum. Importantly, using this approach, we identify the non-cell autonomous effects produced by disrupting *Foxp1* in one cell-type and the molecular compensation that occurs in other populations. Finally, we identify Foxp1-regulated target genes within distinct cell-types and connect these molecular changes to functional and behavioral deficits relevant to phenotypes described in patients with *FOXP1* loss-of-function mutations. These data reveal cell-type-specific transcriptional mechanisms underlying distinct features of striatal circuitry and identify Foxp1 as a key regulator of striatal development.

## Introduction

The striatum is the major input nucleus of the basal ganglia and receives dense glutamatergic inputs from the cortex and thalamus, as well as dopaminergic innervations from the substantia nigra and other neuromodulatory circuits. The principal neurons that receive and integrate this information within the striatum are GABAergic spiny projection neurons (SPNs)^1^. Proper function of striatal circuitry is essential for coordinated motor control, action selection, and reward-based behaviors^2,3^. Dysfunction of this system is implicated across many neurological disorders, including Huntington’s disease, Parkinson’s disease, autism spectrum disorder (ASD), and obsessive-compulsive disorder^4,5^.

Striatal organization has two prominent features: the division of the striatum into distinct neurochemical zones, the striosome and matrix compartments, and the division of SPNs into the direct or indirect projection pathways. Striosome and matrix compartments are enriched for distinct neuropeptides and contribute differentially to striatal connectivity and behavior^4,6–8^. Recent evidence suggests that striosome-matrix compartmentalization is the initial organizational plan during striatal development with distinct intermediate progenitor pools in the lateral ganglionic eminence (LGE) giving rise first to striosome SPNs then matrix SPNs^9^. These progenitor pools then generate either direct and indirect pathway SPNs, which populate both compartments^9^. Direct pathway SPNs express dopamine receptor 1 (D1, dSPNs) and project to the globus pallidus internal (GPi) and substantia nigra (SN). Indirect pathway SPNs express dopamine receptor 2 (D2, iSPNs) and project to the globus pallidus external^1^. Ultimately, these pathways work to bidirectionally modulate excitatory inputs back onto the cortex^1^.

Mature dSPNs and iSPNs have distinct molecular profiles based on expression profiling studies^10–13^, and several transcription factors and chromatin regulators have been identified for both pan-SPN and d/iSPN sub-specification^14–26^. However, a limitation of these previous studies was that non-cell autonomous changes in gene expression were unable to be detected. Moreover, the cellular diversity of the early postnatal striatum, in general, has not been characterized at single-cell resolution. This time point is an important and understudied period of striatal development before excitatory synaptic density onto SPNs markedly increases and where perturbations of cortical-striatal activity can have long lasting effects on SPN spine density and circuit activity^27,28^.

Forkhead-box protein 1 (Foxp1) is a transcription factor with enriched expression in the striatum compared to the rest of the brain^11^. Foxp1 is highly expressed within both SPN populations and loss-of-function *FOXP1* variants are strongly linked to ASD and intellectual disability in humans^29–31^. Expression of Foxp1 begins in the LGE at E12.5 with enrichment in the marginal zone and is maintained throughout striatal development^26,32^. While previous studies have suggested a role for Foxp1 in striatal development^33,34,35^, no study has examined the contribution of Foxp1 to striatal circuit organization in a cell-specific manner.

To ascertain the cell-type specific role of Foxp1, we generated mice with deletion of *Foxp1* from dSPNs, iSPNs, or both populations, and used a combination of single-cell RNA-sequencing (scRNA-seq), whole brain 3D-imaging, and behavioral assays to delineate the contribution of Foxp1 to striatal development and function. We show that Foxp1 is crucial for maintaining the cellular composition of the striatum, especially iSPN specification, and proper formation of the striosome-matrix compartments. We uncover downstream targets regulated by Foxp1 within iSPNs and dSPNs and connect these molecular findings to cell-type-specific deficits in motor and limbic system-associated behaviors, including motor-learning, ultrasonic vocalizations, and fear conditioning. Moreover, we identify the non-cell autonomous effects produced by disruption of one SPN subpopulation and the molecular compensation that occurs. These findings provide an important molecular window into postnatal striatal development and further our understanding of striatal circuits mediating ASD-relevant behavioral phenotypes.

## Results

### Early postnatal scRNA-seq of striatal cells across *Foxp1 cKO* mice

To examine the contribution of Foxp1 to striatal development in a cell-type-specific manner, we generated *Foxp1* conditional knockout (cKO) mice using BAC-transgenic mice driving Cre expression under the D1- or D2-receptor promoters^36^ crossed to *Foxp1*^*flox/flox*^ mice^37–39^ (**Fig. 1a**). Four genotypes were used for downstream analyses: *Drd1-Cre*^*tg/+*^*; Foxp1*^*flox/flox*^ (***Foxp1*^*D1*^**, deletion of *Foxp1* in dSPNs), *Drd2-Cre* ^*tg/+*^*; Foxp1*^*flox/flox*^ (***Foxp1*^*D2*^**, deletion of *Foxp1* in iSPNs), *Drd1-Cre* ^*tg/+*^*; Drd2-Cre* ^*tg/+*^*; Foxp1*^*flox/flox*^ (***Foxp1*^*DD*^**, deletion of *Foxp1* in both d/iSPNs), and *Foxp1*^*flox/flox*^ (***Foxp1*^*CTL*^**). We confirmed that Foxp1 was reduced at both the transcript and protein levels within the striatum (**Fig. 1b-d**). Foxp1 is also reduced in lower-layer cortical neurons expressing *Drd1* (**Supplementary Fig. 1a**).

**Fig. 1.**
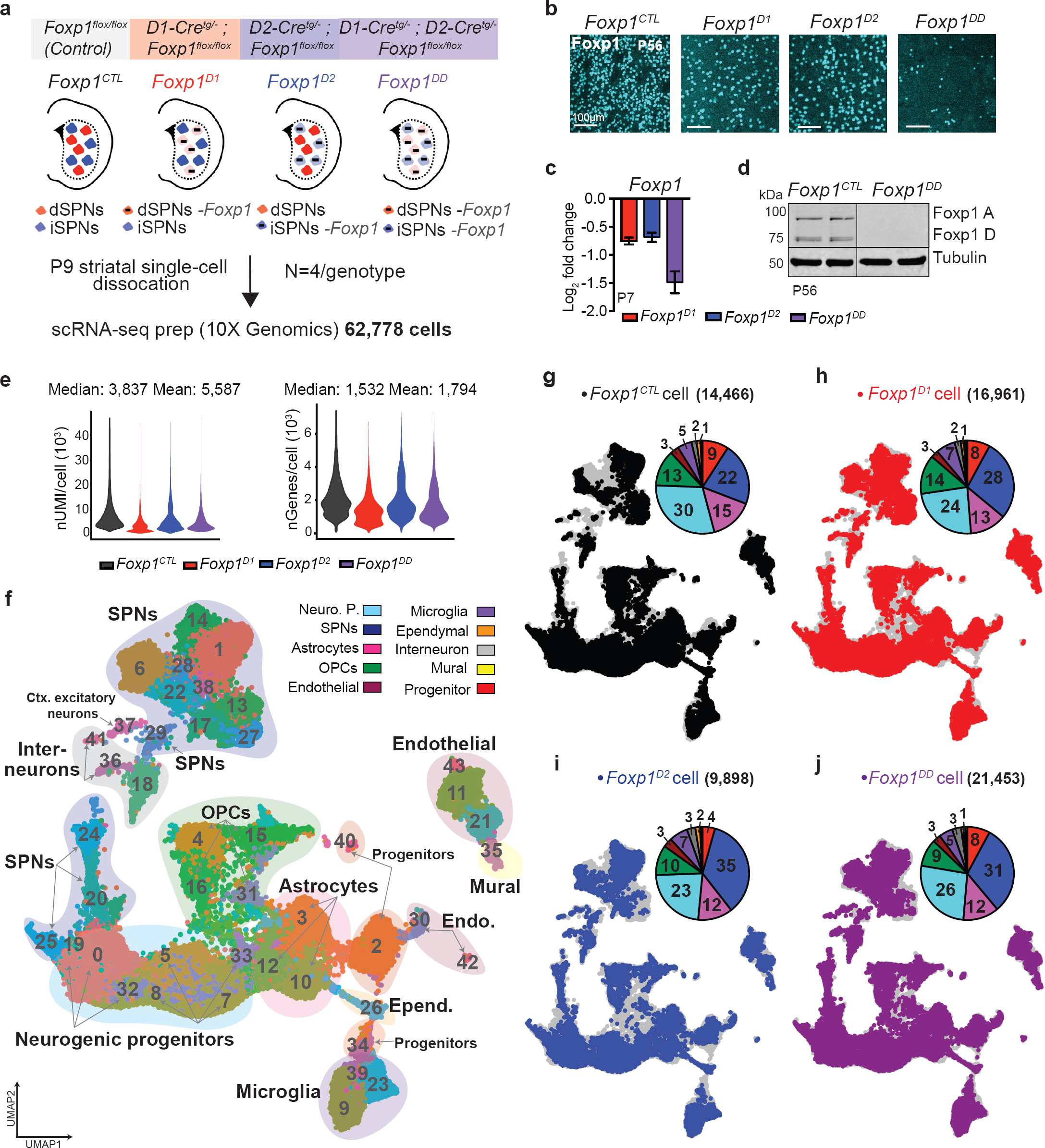
Early postnatal scRNA-seq of striatal cells across *Foxp1 cKOs.* **a**) Schematic of the scRNA-seq experiment using striatal tissue from P9 mice (N=4/genotype) with cell-type-specific conditional deletion of *Foxp1* within the dopamine receptor-1 (*Foxp1*^*D1*^), dopamine receptor-2 (*Foxp1*^*D2*^), or both (*Foxp1*^*DD*^) cell-types. **b**-**d**) Foxp1 is reduced in the striatum via immunohistochemistry (P56) (**b**) and quantitative RT-PCR (P7) (**c**) within each cKO line, with near complete reduction in *Foxp1*^*DD*^ striatal tissue via immunoblot (P56) (**d**) (100μm scale bar). **e**) Violin plots of median and mean number of UMIs or genes per cell across all genotypes. **f**) Non-linear dimensionality reduction with UMAP of all 62,778 post-filtered cells combined across genotype and used for downstream analyses. Cell-type annotation is overlaid to identify the major cell-type represented by each cluster (43 total clusters). **g**-**j**) UMAP plot of cells from (**f**) color-coded to identify each cell by genotype. Pie charts using colors from (**f**) show the striatal cell-type composition as a percentage of total cells within each genotype.

Using 10X Genomics Chromium technology^40^, we profiled the transcriptome of 62,778 striatal cells across control and the three *Foxp1 cKO* mouse lines at postnatal day 9 (N=4/genotype; 16 samples total) (**Fig. 1a**). We detected 5,587 UMIs (median= 3,837) and 1,794 genes (median=1,532) per cell (**Fig. 1e**). All cells were combined across genotype and filtered for downstream clustering, resulting in 43 clusters driven by cell-type (**Fig. 1f** and **Supplementary Table 1**). For unbiased characterization of striatal cell-types, we used a previously annotated adult striatal single-cell dataset^13^ to assign cell-types to each cluster using two separate methods, a previously published expression weighted cell-type enrichment (EWCE) analysis^41^ and an in-house correlation analysis (see methods) (**Supplementary Fig. 1b**). We confirmed the cell-type annotation of our dataset by examining the expression of known marker genes for each major cell-type (**Supplementary Fig. 1c-f** and **Supplementary Table 1**). The principal cell-types found within the early postnatal striatum were neurogenic progenitor cells, spiny projection neurons (SPNs), astrocytes, and oligodendrocyte precursor cells (OPCs) (**Fig. 1f, g** and **Supplementary Table 2**). Endothelial, microglia, ependymal, interneurons, and mural cells made up a smaller percentage of total cells within the postnatal striatum (**Fig. 1f, g**). Unexpectedly, at postnatal day 9, we found a large neurogenic progenitor population (∼30% of total cells within control samples) with clusters expressing proliferation markers (*Mki67*, **Supplementary Fig. 1e**), progenitor markers (*Ascl1, Dlx2*, **Supplementary Fig. 1e**), and SPN-specification markers (*Sp9, Ppp1r1b, Drd1, Drd2*, **Supplementary Fig. 1f**). These data suggest ongoing striatal neuronal differentiation and neurogenesis into early postnatal development.

The cell-type with the largest number of unique subclusters were SPNs with 13 unique clusters (**Fig. 1f**). SPNs and neurogenic progenitors made up 52% of the total cell population (**Fig. 1g**) in line with previously published adult scRNA-seq datasets^12,13^. Genotype-specific variations were observed primarily within SPN clusters, where Foxp1 is selectively deleted (**Fig. 1g-j** and **Supplementary Table 2).** To more directly compare the composition of striatal cell-types across genotypes and better control for variations in total cells sequenced between genotypes, we down-sampled the dataset to yield equal cell numbers across genotypes and reclustered the resultant cells separately. We found analogous results in the percentage of cell-types from down-sampling experiments compared to the full dataset (**Supplementary Fig. 1g**). Variations within the SPN and progenitor populations in *Foxp1 cKO* samples compared to control were consistent across down-sampling, with more neurons (SPNs) and fewer progenitor cells within all *Foxp1 cKO* samples (**Fig. 1g-j, Supplementary Fig. 1g**, and **Supplementary Table 2**). Our data reveal at the single-cell level that deletion of *Foxp1* reduces the population of striatal neurogenic cells, while increasing the percentage of mature SPNs. These data highlight the diversity of the cellular composition of the early-postnatal striatum and demonstrate that Foxp1 plays an important role in striatal neurogenesis and development.

### Diversity of early post-natal striatal projection neurons

To further characterize early postnatal SPN subtypes and the effects of *Foxp1* deletion, we next isolated all clusters identified as neuronal from the annotation analyses (see Figure 1 and methods) and reclustered them separately (18,073 cells total and 24 clusters) (**Fig. 2a-e**). Three interneuron clusters (Clusters-14, 15, 20) were identified by the interneuron marker *Nkx2*-*1* (**Fig. 2a, Supplementary Fig. 2a**, and **Supplementary Table 1**). We could clearly distinguish dSPN clusters (Clusters-0, 1, 3, 4, 5, 9) and iSPN clusters (Clusters-2, 8, 10, 16) using canonical markers (*Drd1* and *Tac1* for dSPNs, *Drd2* and *Penk* for iSPNs) (**Fig. 2f, Supplementary Fig. 2b**, and **Supplementary Table 1**). Pairwise comparisons between the major dSPN and iSPN clusters confirmed enrichment of known genes within each population (**Supplementary Fig. 2c**). One small cluster (Cluster-19) co-expressed both *Drd1* and *Drd2* receptors, termed “ddSPNs” (**Fig. 2f**). SPNs expressing both *Drd1* and *Drd2* receptors were also scattered throughout other clusters and comprised ∼1% of the total SPN population (**Supplementary Fig. 2d** and **Supplementary Table 2**). We identified a recently described SPN subpopulation termed “eccentric” SPNs (eSPNs)^13^ within Cluster-7 that uniquely expressed markers such as *Casz1* and *Otof* (**Fig. 2f, Supplementary Fig. 2e**, and **Supplementary Table 1**). We also found two clusters (Cluster-6, 23) that were enriched for the neurogenic transcription factors *Sox4* (**Fig. 2f** and **Supplementary Fig. 2f**) and *Sox11* (**Supplementary Table 1**). *Sox4* and *Sox11* function during the terminal steps of neurogenesis to promote neuronal maturation^42,43^; therefore, we termed these clusters “immature SPNs” (imSPNs). We confirmed the presence of Sox4^+^ cells within and near the subventricular zone of the lateral ventricle and populating zones in P7 ventral striatum (**Supplementary Fig. 2g**). Additionally, several clusters enriched for d/iSPN markers also have high expression of *Sox4* (dSPN Clusters-9,11, 13, 17 and iSPN Clusters-16), indicating these may be less mature SPNs (**Fig. 2g** and **Supplementary Fig. 2f**). Two clusters (Cluster-12, 18) were composed primarily of cells from *Foxp1*^*D1*^ and *Foxp1*^*D2*^ and could not be classified into distinct SPN subclusters (**Fig. 2a-f)**. *Foxp2*, another Foxp transcription factor with high amino acid sequence similarity to *Foxp1*^44^, is an SPN marker with enriched expression in dSPNs (**Supplementary Fig. 2c**)^29,45^. Within our dataset, *Foxp2* is highly expressed within all dSPN clusters and one iSPN cluster (Cluster-8). Surprisingly, the highest expression of *Foxp2* is found within eSPN Cluster-7 and imSPN Cluster-6, where notably *Foxp1* is not highly expressed (**Fig. 2f** and **Supplementary Fig. 2h**). *Foxp2* expression is also maintained within adult eSPNs^13^. We confirmed that Foxp2 is expressed in cells other than mature dSPNs and iSPNs at postnatal day 9 using *D1-tdTomato*^*tg/-*^ and *D2-eGFP*^*tg/-*^ reporter mice (**Supplementary Fig. 2i**).

**Fig. 2.**
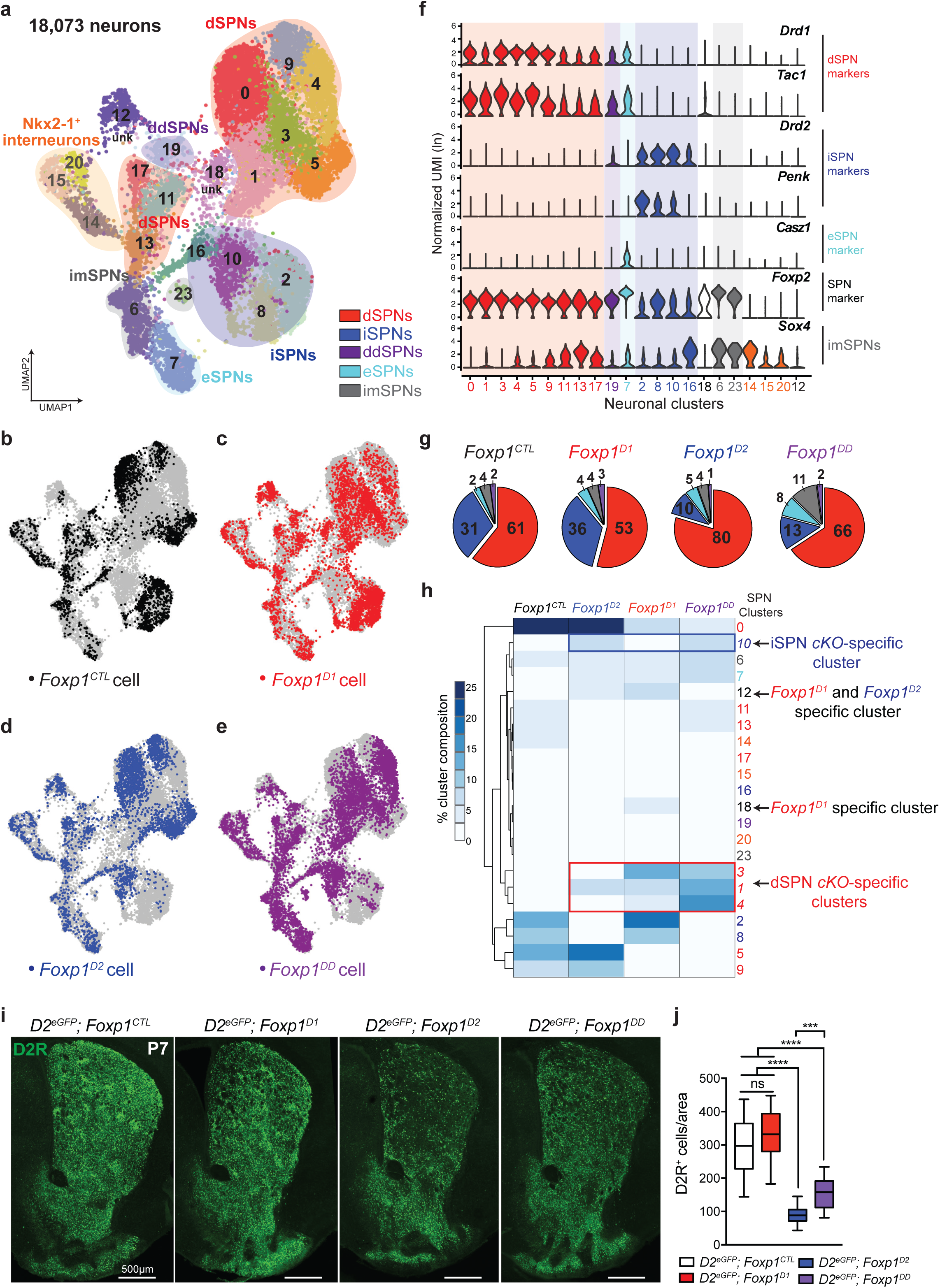
Foxp1 specifies distinct SPN subpopulations. **a**) UMAP plot showing each neuronal subcluster by color with overlay colors showing neuronal subpopulation identity. **b**-**e**) UMAP plots of cells from (**a**) color-coded to identify each cell by genotype. **f**) Violin plots of the normalized UMI expression of markers of SPN subpopulations: dSPNs (*Drd1, Tac1, Foxp2*), iSPNs (*Drd2, Penk*), ddSPNs (*Drd2, Drd1, Tac1*), eSPNs (*Casz1*), and imSPNs (*Sox4*). **g**) Pie charts showing altered composition of SPN subtypes within *Foxp1 cKO* mice (using colors from **a**). **h**) Heatmap showing the percentage of cells contributing to each cluster across genotype (using colors from **a**). **i**-**j**) *Foxp1 cKO* mice were crossed to *D2*^*eGFP*^ reporter lines to label dopamine receptor-2 (D2R) iSPNs in green (coronal section, 500μm scale bar). *Foxp1*^*D2*^ *and Foxp1*^*DD*^ mice had significantly fewer iSPNs compared to *Foxp1*^*CTL*^ mice at P7, while *Foxp1*^*DD*^ mice had significantly more iSPNs compared to *Foxp1*^*D2*^ animals. Data are represented as a box plot, n=3-6 mice/genotype. *\*\*\*\**p*<*0.0001, *\*\*\**p*<*0.005, one-way ANOVA with Tukey’s multiple comparisons test.

### Foxp1 regulates SPN subtype composition and iSPN specification

We next asked whether Foxp1 regulates the development of specific SPN populations by examining the percentages of SPN subtypes across genotypes (**Fig. 2g**). Control samples have nearly double the number of dSPN relative to iSPNs (61% dSPNs, 31% iSPNs), with imSPNs contributing ∼4% of the total SPN population and both eSPNs and ddSPNs contributing ∼2% (**Fig. 2g**). This percentage of dSPNs to iSPNs at P9 is similar to those seen at P14 using reporter mice^46^. The percentage of SPN subtypes varied across *Foxp1 cKO* samples (**Fig. 2g**). Notably, the number of eSPNs increased 2-4-fold across *Foxp1 cKO* samples. Strikingly, within *Foxp1*^*D2*^ and *Foxp1*^*DD*^ samples, the number of iSPNs was reduced by two-thirds compared to control levels (**Fig. 2g**). Cells with deletion of *Foxp1* were transcriptionally distinct and clustered largely separately from control cells (**Fig. 2h**).

To independently confirm the reduction of iSPNs in *Foxp1*^*D2*^ and *Foxp1*^*DD*^ samples, we crossed all *Foxp1 cKO* mice to *D2-eGFP* reporter animals (*D2-eGFP*^*tg/-*^*; Foxp1*^*flox/flox*^) to label iSPNs (**Fig. 2i**). Within *Foxp1*^*D2*^ mice, we again found a significant two-thirds reduction of iSPNs (D2-eGFP^+^ cells) as seen in the scRNA-seq data (**Fig. 2i, j**). Compared to *Foxp1*^*CTL*^, *Foxp1*^*DD*^ mice also showed significantly reduced iSPNs, but they also showed increased iSPNs compared to *Foxp1*^*D2*^ (**Fig. 2i, j**). The remaining iSPNs in the *Foxp1*^*D2*^ mice were not the product of *D2-Cre* inefficiency, as these cells did not express Foxp1 (**Supplementary Fig. 2j**). Only 7 iSPNs within the single-cell *Foxp1*^*CTL*^ data did not express Foxp1 (0.2% of total iSPNs) (**Supplementary Fig. 2d**), therefore, we would not expect the remaining iSPNs in *Foxp1*^*D2*^ mice to be a naturally occurring Foxp1 negative population. Taken together, these results indicate that Foxp1 is required for the specification of distinct iSPN subpopulations and may function to repress the generation of eSPNs.

### Deletion of *Foxp1* disrupts striosomal area and iSPN localization

We identified distinct subclusters within dSPNs and iSPNs in our scRNA-seq data (**Fig. 3a**). Using a pairwise differential gene expression analysis between clusters, we found that two *Foxp1*^*CTL*^ dSPN clusters (Cluster-0, 5) corresponded to either matrix or striosome compartments, respectively, based on the enrichment of known striosome (*Oprm1, Isl1, Pdyn, Lypd1, Tac1*, and *Nnat*) or matrix markers (*Ebf1, Epha4, Mef2c*) (**Fig. 3b** and **Supplementary Table 3**). Within iSPNs, Cluster-8 was enriched for striosomal markers (*Nnat, Lypd1, Foxp2*) and Cluster-2 for matrix markers (*Penk, Chrm3, Epha4*) (**Fig. 3c** and **Supplementary Table 3**).

**Fig. 3.**
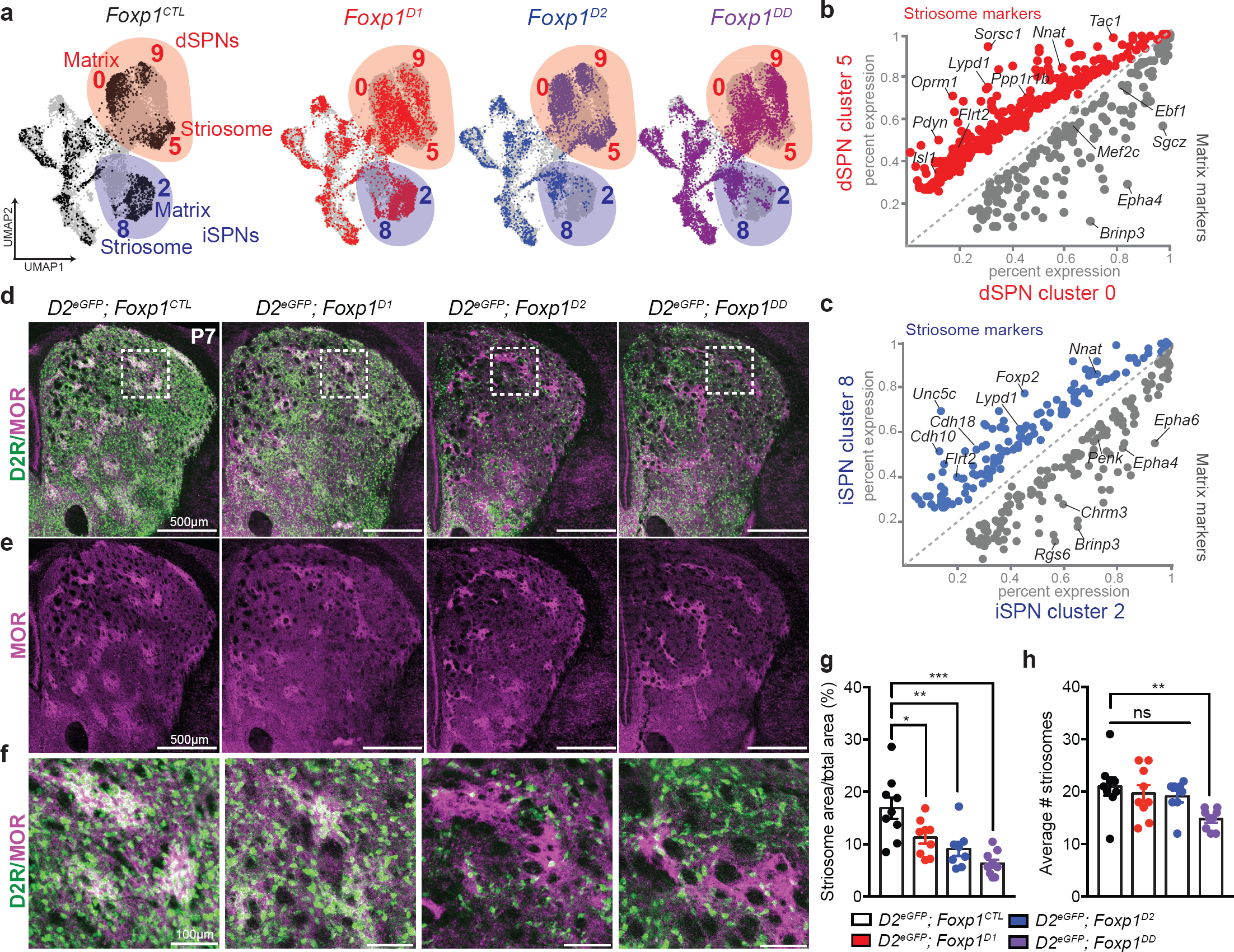
Foxp1 regulates striosome-matrix organization. **a**) Within *Foxp1*^*CTL*^ samples, dSPN and iSPNs have large sub-clusters (Clusters-0 and -5 for dSPNs and Clusters-2 and -8 for iSPNs). Cells with deletion of *Foxp1* cluster largely separately from control cells and subclusters within iSPNs and dSPNs are more intermixed (*Foxp1*^*D1*^ dSPNs) or lost completely (*Foxp1*^*D2*^ iSPNs). **b**-**c**) Scatter plots showing the percent expression of enriched transcripts between Clusters-0 and-5 (**b**) or Clusters-2 and-8 (**c**). Striosome markers are enriched in dSPN Cluster-5 and iSPN Cluster-8, while matrix markers are enriched in dSPN Cluster-0 and iSPN Cluster-2 (p.adj<0.05). **d**-**f**) iSPNs within *Foxp1*^*D2*^ and *Foxp1*^*DD*^ mice localized primarily along the striosomal border marked by IHC for Mu-Opiod Receptor (MOR) in P7 animals crossed to D2-eGFP reporter mice (500μm scale bar in D-E, 100μm scale bar in **f**). **g**-**h**) The striosome compartment was significantly reduced across all *Foxp1 cKO* mice as a percent of total striatal area (measuring only dorsal striosomes) and the number of striosome “patches” was significantly reduced in *Foxp1*^*DD*^ animals. Data are represented as mean ± SEM, n=4 mice/genotype. *\**p<0.05, p**<0.005, *\*\*\**p*<*0.0001, one-way ANOVA with Tukey’s multiple comparisons test.

We next wanted to determine whether the remaining subpopulation of iSPNs within *Foxp1*^*D2*^ or *Foxp1*^*DD*^ mice localized within either the striosome or matrix compartment. To do this, we stained for the canonical striosome marker MOR (*Oprm1*) in *Foxp1 cKO* mice crossed to *D2-eGFP* reporter mice (**Fig. 3d-f**). We found that few remaining iSPNs within *Foxp1*^*D2*^ and *Foxp1*^*DD*^ mice localized within the striosome compartment. They clustered primarily around the border of the striosome compartments and were scattered throughout the matrix (**Fig. 3d-f**). Taken together, these data show that Foxp1 specifies iSPNs of both striosome and matrix compartments but may not be necessary for specification of a subpopulation of iSPNs near the striosomal border. We also found that striosomal area was significantly reduced across all *Foxp1 cKO* animals at P7 (**Fig. 3g**) and that fewer striosome “patches” were observed specifically within *Foxp1*^*DD*^ mice (**Fig. 3h**). These data indicate that Foxp1 plays a critical role within both dSPNs and iSPNs to maintain proper striosome-matrix architecture.

### Cell-type-specific Foxp1 regulated targets

To better understand the molecular mechanisms regulated by Foxp1, we performed a cell-type-specific “pseudobulk” differential gene expression analysis (see methods) of the scRNA-seq data across genotypes. We identified differentially expressed genes (DEGs) regulated by *Foxp1* within dSPNs or iSPNs, both cell-autonomously and non-cell-autonomously (**Fig. 4a, b** and **Supplementary Table 4**). Cell-autonomous DEGs are found in Cre active cells (dSPNs in *Foxp1*^*D1*^ samples or iSPNs in *Foxp1*^*D2*^ samples) and non-cell-autonomous DEGs are found in Cre inactive cells (iSPNs in *Foxp1*^*D1*^ samples or dSPNs in *Foxp1*^*D2*^ samples). We observed more total iSPN-DEGs (647) compared to dSPNs-DEGs (285) across genotypes (**Fig. 4a, b**). There were more cell-autonomous changes than non-cell-autonomous within both dSPNs and iSPNs of *Foxp1*^*D1*^ and *Foxp1*^*D2*^ samples and no differences in the ratio of cell-autonomous to non-cell autonomous DEGs within dSPNs or iSPNs were observed (**Fig. 4c**). However, significantly more iSPN-DEGs were shared between *Foxp1*^*D2*^ and *Foxp1*^*DD*^ samples (211 DEGs) compared to dSPN-DEGs shared between *Foxp1*^*D1*^ and *Foxp1*^*DD*^ samples (47 DEGs) (**Fig. 4d**). The DEGs unique to *Foxp1*^*DD*^ samples were termed “interaction-DEGs”. We found significantly more interaction-DEGs in dSPNs suggesting that iSPN dysfunction exerts more transcriptional changes upon dSPNs than vice versa (**Fig. 4d**).

**Fig. 4.**
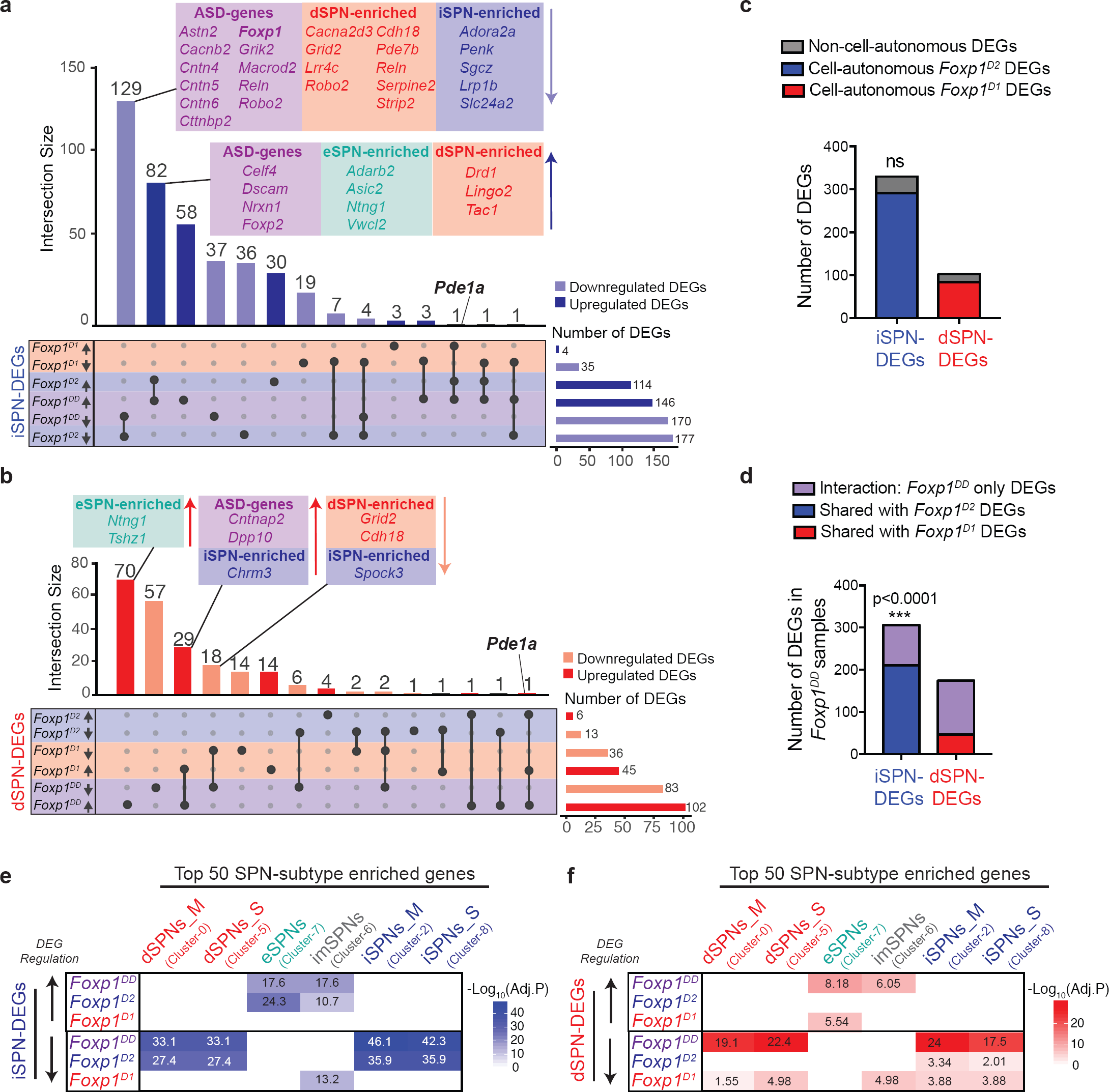
Foxp1 regulates cell-type-specific molecular pathways. **a**-**b**) SPN cell-type-specific differential gene expression between genotypes. Upset plot showing the overlap of upregulated or downregulated DEGs across genotypes within iSPNs (**a**) or dSPNs (**b**). Genes shown within boxes are color-coded by categories indicated. **c**) No significant difference between the number of DEGs within iSPNs and dSPNs that are cell-autonomous vs non-cell-autonomous (p=0.0975, two-sided Fisher’s exact test). **d**) There is a significant difference in the number of DEGs within *Foxp1*^*DD*^ mice that overlap with *Foxp1*^*D2*^ or *Foxp1*^*D1*^ DEGs to unique *Foxp1*^*DD*^ DEGs (interaction DEGs) (p<0.0001, two-sided Fisher’s exact test). **e**-**f**) Enrichment of upregulated or downregulated iSPN-DEGs (**e**) or dSPN-DEGs (**f**) across *Foxp1 cKO* samples in distinct SPN subtypes (top 50 most enriched genes/cluster) using a hypergeometric overlap test (8,000 genes used as background).

The striking difference in total number of DEGs between iSPNs and dSPNs could be due to transcriptional compensation by Foxp2 in dSPNs. Foxp2 is enriched in dSPNs relative to iSPNs (**Supplementary Fig. 2c**) and we previously found that Foxp1 and Foxp2 have shared striatal targets^35^. Interestingly, *Foxp2* is increased in iSPNs with loss of Foxp1, suggesting that Foxp1 may function to repress Foxp2 within distinct iSPN subtypes (**Fig. 4a** and **Supplementary Table 4**). *Six3* (Six homeobox 3), a transcription factor crucial for iSPN specification^26^, is also upregulated within the remaining iSPNs of *Foxp1*^*D2*^ and *Foxp1*^*DD*^ mice (**Supplementary Table 4**). We previously found that *SIX3* was a direct target of FOXP1 in human neural progenitors^35^. Therefore, upregulation of both *Foxp2* and *Six3* in iSPNs may play a role in the specification of the remaining iSPNs within *Foxp1*^*D2*^ and *Foxp1*^*DD*^ mice.

Gene ontology (GO) analysis of the shared iSPN-DEGs within *Foxp1*^*D2*^ and *Foxp1*^*DD*^ supports a role for Foxp1 in axon guidance, neurogenesis, and neuronal differentiation of iSPNs (**Supplementary Table 5**). Shared upregulated dSPN-DEGs within *Foxp1*^*D1*^ and *Foxp1*^*DD*^ suggest altered synaptic and voltage-gated mechanisms (**Supplementary Table 5**). We confirmed changes in cell-type-specific gene expression via immunohistochemistry for a subset of top DEGs (*Pde1a, Calb1*, and *Darpp32*) using dual-reporter mice labelling dSPNs with tdTomato (*Drd1-tdTomato*^*tg/+*^*; Foxp1*^*flox/flox*^) and iSPNs with eGFP (*Drd2-eGFP*^*tg/+*^*; Foxp1*^*flox/flox*^) crossed to *Foxp1 cKO* strains (**Supplementary Fig. 3a-e**). *Pde1a*, a gene encoding a calmodulin/Ca^2+^ activated phosphodiesterase, was upregulated in both SPN subtypes within all *Foxp1 cKO* samples in a cell autonomous and non-cell-autonomous manner (**Fig. 4a, b and Supplementary Fig. 3a, d-e**). Previous *in vitro* work found that loss of Foxp1 reduced the expression of DARPP-32 (*Ppp1r1b*), a critical phosphatase in the dopamine signaling cascade^33^. We show this decrease in DARPP-32 is specific to iSPNs *in vivo* (**Supplementary Fig. 3b, d-e**). We also confirmed the increase of calbindin 1 (*Calb1*) selectively in dSPNs with deletion of *Foxp1* (**Supplementary Fig. 3c-e**).

Given our previous finding that striatal targets of Foxp1 overlapped significantly with ASD-associated genes^35^, we examined the cell-type-specificity of this overlap (**Supplementary Fig. 3f**). Using the SFARI ASD gene list, we found a significant overlap with high-confidence ASD-risk genes (SFARI gene score 1-4) with iSPNs-DEGs with cell-autonomous deletion of *Foxp1*. These genes included three members of the contactin-family of axon-associated cell adhesion molecules: *Cntn4, Cntn5, Cntn6* (**Fig. 4a, Supplementary Fig. 3f**). There was no significant overlap with ASD-risk genes and cell-autonomous DEGs in dSPNs (**Supplementary Fig. 3f**). Surprisingly, we found a significant overlap with upregulated, non-cell autonomous iSPN-DEGs within *Foxp1*^*D1*^ samples (*Kirrel3, Nlgn1*) (**Supplementary Fig. 3f**). Both iSPN- and dSPN-DEGs within *Foxp1*^*DD*^ samples overlapped with ASD-risk genes (**Supplementary Fig. 3f**). These data demonstrate that cell-type-specific deletion of *Foxp1* specifically within iSPNs modulates ASD-associated molecular pathways both cell-autonomously and non-cell-autonomously.

Two ASD-risk genes that were upregulated with deletion of *Foxp1* in dSPNs were *Cntnap2* (contactin-associated protein like 2) and *Dpp10* (dipeptidyl peptidase like 10) (**Fig. 4b** and **Supplementary Table 4**). *Cntnap2* is a known repressed downstream target of both Foxp1 and Foxp2^47,48^ and we previously found upregulation of *Dpp10* within *Foxp1*^*+/-*^ striatal tissue using bulk RNA-sequencing ^35^. Here, using scRNA-seq, we show this regulation is specific to dSPNs.

### Upregulation of eSPN molecular markers with deletion of *Foxp1*

To determine whether deletion of *Foxp1* within SPNs altered cell identity, we overlapped the top 50 enriched gene markers of distinct SPN subpopulations (eSPNs, imSPNs, and matrix and striosome dSPNs and iSPNs) (**Supplementary Table 1**) with upregulated or downregulated iSPN-DEGs (**Fig. 4e**) or dSPN-DEGs (**Fig. 4f**) found within each *Foxp1 cKO* group. The upregulated DEGs in both iSPNs and dSPNs with cell-autonomous deletion of *Foxp1* were significantly enriched for molecular markers of eSPNs. Upregulated iSPN-DEGs were specifically enriched for the top four enriched eSPNs markers: *Adarb2, Ntng1, Asic2*, and *Foxp2* (**Fig. 4a, e-f**). iSPN and dSPN subtype enriched genes significantly overlapped with downregulated DEGs in both *Foxp1*^*D1*^ and *Foxp1*^*D2*^ samples (**Fig. 4a, e-f**). Taken together, these results indicate that Foxp1 is important for maintaining the molecular identity of dSPNs and iSPNs within both matrix and striosome compartments and repressing eSPN molecular identity within these cell-types.

### Altered direct and indirect pathway projections in *Foxp1*^*D2*^ mice

Many DEGs regulated by Foxp1 within SPNs are involved in axonogenesis and neuron projection (**Supplementary Tables 4 and 5**). We therefore examined SPN projection patterns impacted by cell-type-specific deletion of *Foxp1* in adult mice using serial two-photon tomography combined with a machine-learning-based quantification algorithm^49,50^. We crossed *Foxp1*^*D1*^ *and Foxp1*^*D2*^ mice to D1-tdTomato and/or D2eGFP reporter mice (described above) to visualize projection patterns of both the direct (dSPN) and indirect (iSPN) pathway, respectively. We first quantified total striatal area across genotypes and found a significant decrease in striatal area in *Foxp1*^*D2*^ mice, while no changes were found in *Foxp1*^*D1*^ animals (**Fig. 5a**). We next found a significant reduction of iSPN terminals onto the GPe in *Foxp1*^*D2*^ mice, which was not unexpected given the significant decrease in iSPNs (**Fig. 5b, d-e**). iSPN terminals onto the GPe were unaltered in *Foxp1*^*D1*^ mice (**Fig. 5b-c, e**). Moreover, there were no changes in dSPN projection patterns in *Foxp1*^*D1*^ mice; however, *Foxp1*^*D2*^ mice had significant deficits in dSPN projections onto the GPi, supporting a non-cell-autonomous role for Foxp1 in iSPNs (**Fig. 5b-d, f**). These findings indicate that Foxp1 regulates both iSPN and dSPN projection patterns through its role in iSPNs (**Fig. 5g**). Within our scRNA-seq data, non-cell-autonomous dSPN-DEGs in *Foxp1*^*D2*^ samples were enriched for GO categories such as neuron projection (**Supplementary Table 5**). Since projections onto the GPi were not altered in *Foxp1*^*D1*^ mice, dSPN-DEGs unique to *Foxp1*^*D2*^ samples are most likely responsible for the altered dSPN projection patterns found within *Foxp1*^*D2*^ animals. We therefore examined the overlap of dSPN-DEGs within *Foxp1*^*D1*^ (cell-autonomous) and *Foxp1*^*D2*^ samples (non-cell-autonomous) (**Fig. 5h**). dSPN-DEGs unique to *Foxp1*^*D2*^ samples involved in neuron projection include *Akap5, Asic2, Kirrel3, Cdh8*, and *Cntn4* (**Fig. 5h**). Interestingly, *Kirrel3, Cdh8*, and *Cntn4* are also ASD-risk genes (**Fig. 5h**). These findings suggest deletion of *Foxp1* within iSPNs alters the gene expression profiles within both iSPNs and dSPNs important for proper striatal projection patterning.

**Fig. 5.**
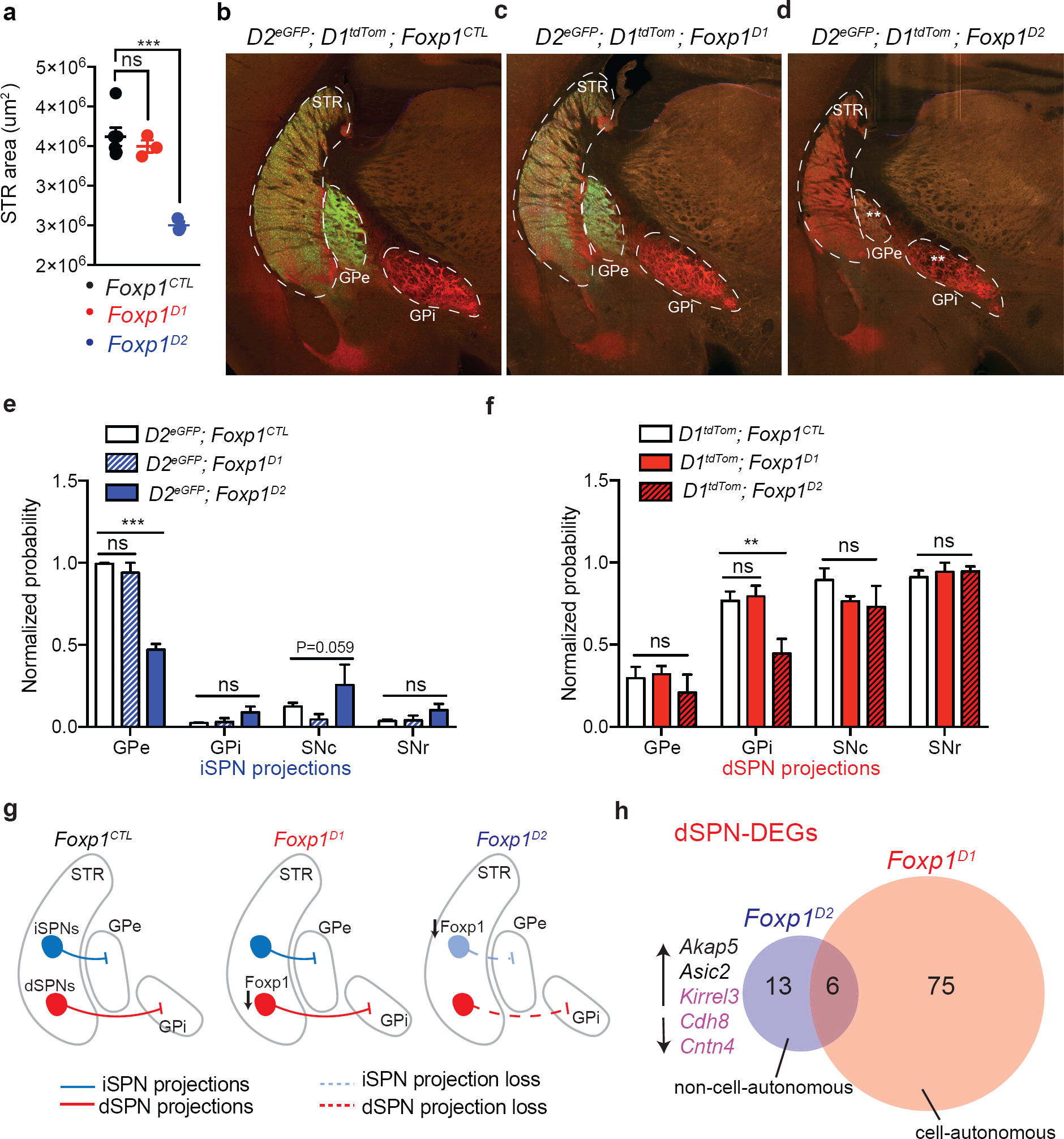
Deletion of *Foxp1* in iSPNs alters projection patterns of both dSPNs and iSPNs. **a**) Striatal area quantification of four serial slices from anterior to posterior at 400um increments within *Foxp1*^*CTL*^, *Foxp1*^*D1*^, and *Foxp1*^*D2*^ adult mice. Data are represented as mean ± SEM, n=3-4 mice/genotype. *\*\*\**p*<*0.001, one-way ANOVA with Tukey’s multiple comparisons test. **b**-**d**) Representative Tissuecyte 1000 coronal section showing the projections of dSPNs and iSPNs using *D1*^*tdTom*^ and *D2*^*eGFP*^ reporter mice, respectively, crossed to *Foxp1*^*CTL*^ (**b**), *Foxp1*^*D1*^ (**c**), or *Foxp1*^*D2*^(**d**). **e**) Quantification of the normalized probability maps of iSPN (eGFP) projections within *Foxp1*^*CTL*^, *Foxp1*^*D1*^, and *Foxp1*^*D2*^ mice showing reduced GPe projections from iSPNs within *Foxp1*^*D2*^ mice. No significant changes were seen in projection patterns onto the SNc or SNr. Data are represented as mean ± SEM, n=3-4 mice/genotype. *\*\*\**p*<*0.0001, two-way ANOVA with Dunnett’s multiple comparisons test. **f**) Quantification of the normalized probability maps of dSPN (tdTomato) projections within *Foxp1*^*CTL*^, *Foxp1*^*D1*^, and *Foxp1*^*D2*^ mice showing reduced GPi projections from dSPNs within *Foxp1*^*D2*^ mice. Data are represented as mean ± SEM, n=2-4 mice/genotype. *\*\**p*<*0.01, two-way ANOVA with Dunnett’s multiple comparisons test. **g**) Schematic of cell-autonomous and non-cell-autonomous projection deficits found in the *Foxp1*^*D2*^ animals. **h**) Overlap of dSPN-DEGs within *Foxp1*^*D1*^ or *Foxp1*^*D2*^ cells. Unique *Foxp1*^*D2*^ dSPN-DEGs that are involved in neuron projection are shown, with ASD-risk genes highlighted in purple. GPi= globus pallidus internal, GPe= globus pallidus external, STR= striatum, SNr=substantia nigra pars reticulata, SNc= substantia nigra pars compacta.

### Distinct behavioral deficits with cell-type-specific deletion of *Foxp1*

We hypothesized that severe reduction of iSPNs and altered projection patterns with deletion of *Foxp1* from iSPNs would result in altered motor behaviors. We therefore first tested behaviors classically characterized as being governed by striatal circuits, such as motor learning and activity levels. To test motor learning, we used the accelerating rotarod assay and found that *Foxp1*^*D2*^ and *Foxp1*^*DD*^ mice had significant deficits at remaining on the accelerating beam compared to control and *Foxp1*^*D1*^ mice (**Fig. 6a**). This phenotype was not due to differences in grip strength (**Supplementary Fig. 4a, b**) or gait abnormalities (**Supplementary Fig. 4c-f**). *Foxp1*^*D2*^ and *Foxp1*^*DD*^ mice were also hyperactive in the open field behavioral paradigm compared to control mice **(Fig. 6b**); however, no difference was observed in novel cage activity between genotypes (**Supplementary Fig. 4g**). There was no difference in time spent in the periphery versus the center of the open field between genotypes (**Fig. 6c**), suggesting no changes in anxiety-like behavior.

**Fig. 6.**
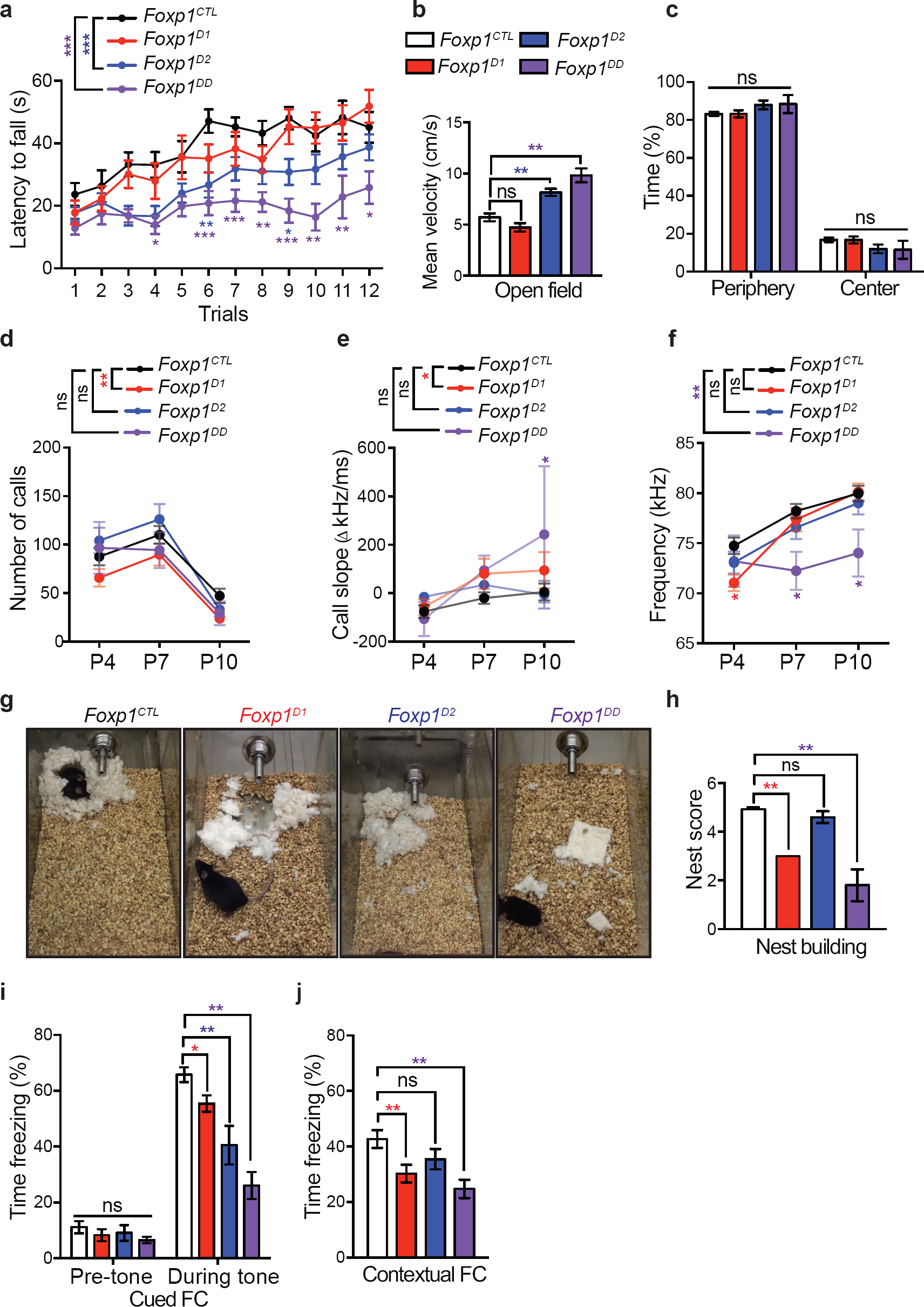
Foxp1 regulates behaviors via distinct striatal circuits. **a**) Latency to fall was measured on the accelerating rotarod. *Foxp1*^*D2*^ and *Foxp1*^*DD*^ mice exhibit significant deficits. Data are represented as mean ± SEM, n=11 *Foxp1*^*CTL*^; n=17 *Foxp1*^*D1*^; n= 18 *Foxp1*^*D2*^; n=12 *Foxp1*^*DD*^. *p<0.05, **p<0.005, ***p<0.0001, two-way ANOVA with Sidak’s multiple comparisons test. **b**-**c**) Mice were tested within the open field paradigm with velocity (**b**) and percent time spent in the periphery vs center (**c**) plotted. *Foxp1*^*D2*^ and *Foxp1*^*DD*^ mice had significant increase in activity with no difference in percent time spent in the periphery and center. Data are represented as mean ± SEM. n=4 *Foxp1*^*DD*^; n=14 *Foxp1*^*D1*^; n=17 *Foxp1*^*D2*^; n=22 *Foxp1*^*CTL*^. *\*\*\**p*<*0.0001, one-way ANOVA with Sidak’s multiple comparisons test. **d**-**f**) Neonatal isolation vocalizations were measured at P4, P7, and P10. (**d**) The number of isolation calls were significantly reduced in *Foxp1*^*D1*^ mice. (**e**) Mean frequency (kHz) of the isolation calls was significantly altered in *Foxp1*^*DD*^ mice and at P4 within *Foxp1*^*D1*^ animals. (**f**) The call slope or “structure” of the call was significantly altered over postnatal development in *Foxp1*^*D1*^ pups and specifically at P10 within *Foxp1*^*DD*^ pups. Data are represented as mean ± SEM. n=11 *Foxp1*^*DD*^; n=47 *Foxp1*^*D1*^; n=36 *Foxp1*^*D2*^; n=71 *Foxp1*^*CTL*^. *p<0.05, **p<0.005, ***p<0.0001, two-way ANOVA with Sidak’s multiple comparisons test. **g**) Representative images of nests. **h**) *Foxp1*^*D1*^ and *Foxp1*^*DD*^ mice produced nests with significantly lower quality scores compared to *Foxp1*^*D2*^ and *Foxp1*^*DD*^ mice. Data are represented as mean ± SEM. n=5 *Foxp1*^*DD*^; n=4 *Foxp1*^*D1*^; n=5 *Foxp1*^*D2*^; n=7 *Foxp1*^*CTL*^. **p<0.005, one-way ANOVA with Sidak’s multiple comparisons test. **i**-**j**) Associative fear memory was assessed using the fear conditioning (FC) paradigm. All *Foxp1 cKO* mice displays deficits in cued FC (**h**) shown as the percent of time spent freezing. Only *Foxp1*^*D1*^ and *Foxp1*^*DD*^ mice displayed deficits in contextual FC (**i**). Data are represented as mean ± SEM. n=15 *Foxp1*^*DD*^; n=22 *Foxp1*^*D1*^; n=11 *Foxp1*^*D2*^; n=23 *Foxp1*^*CTL*^. *p<0.05, **p<0.005, ***p<0.0001, two-way ANOVA with Dunnett’s multiple comparisons test.

Since genetic variants in *FOXP1* are strongly associated with ASD, we next examined ASD-relevant social communication behaviors. Using a maternal separation paradigm, we recorded pup ultrasonic vocalizations (USVs) at three postnatal time points (P4, P7, and P10). We found that *Foxp1*^*D1*^ mice produced significantly fewer calls with altered call slope compared to control pups (**Fig. 6d-e**). In addition, *Foxp1*^*D1*^ pups had significantly lower pitch at P4, while *Foxp1*^*DD*^ mice exhibited deficits in pitch across all developmental time points (**Fig. 6f**). No significant USV changes were measured solely in *Foxp1*^*D2*^ pups. We also tested nest building behavior, an important communal behavior in rodents^51,52^, and found that *Foxp1*^*D1*^ and *Foxp1*^*DD*^ mice produced low-quality nests compared to control and *Foxp2*^*D2*^ nests (**Fig. 6g-h**).

Because individuals with *FOXP1* mutations are frequently comorbid for intellectual disability^30,31^, we next assessed whether learning and memory circuits were altered using the cued and contextual fear conditioning (FC) paradigm (**Fig. 6i-j**). All *Foxp1 cKO* mice had significantly reduced freezing behavior during cued-evoked fear memory recall (**Fig. 6i**); however, only *Foxp1*^*D1*^ and *Foxp1*^*DD*^ mice showed significant deficits in context-evoked fear memory (**Fig. 6j**). While hippocampal and amygdala circuits are classically associated with fear conditioning, striatal D1 receptors are also important for mediating proper contextual FC in mice^53^. We also found that striosome-matrix architecture was more severely disrupted over postnatal development in *Foxp1*^*D1*^ and *Foxp1*^*DD*^ adult animals compared to control and *Foxp1*^*D2*^ mice (**Supplementary Fig. 4h**).

## Discussion

In this study, we use single-cell transcriptomics to examine the molecular mechanisms underlying striatal neuronal specification by sequencing thousands of striatal cells across control and cell-type-specific *Foxp1* conditional mouse models. We show that Foxp1 influences striatal development through cell-type-specific molecular pathways and describe the molecular, functional, and behavioral consequences of *Foxp1* deletion within distinct striatal circuits (**Fig. 7**).

**Figure 7.**
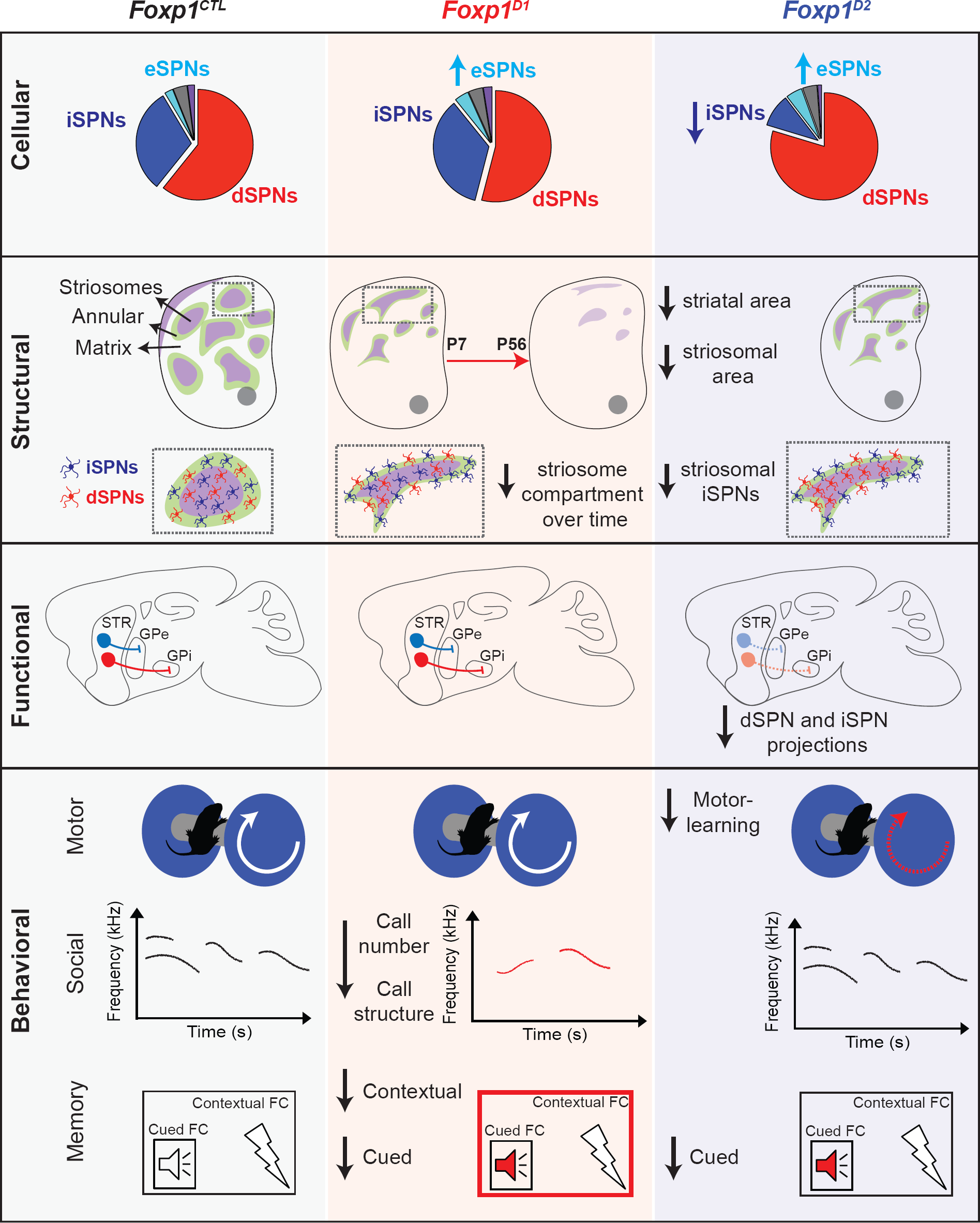
Summary of cellular, structural, functional, and behavioral findings within cell-type-specific *Foxp1* conditional knockout mice. *Foxp1*^*D1*^ mice have an increase in eSPN subpopulations, reduced striosomal area, no gross SPN projection deficits, and distinct behavioral deficits relevant to social communication behavior and contextual fear conditioning. *Foxp1*^*D2*^ mice have a marked decrease in iSPN and increase in eSPN subpopulations, reduced striosomal area with few striosomal iSPNs, dSPN and iSPN projection deficits, and distinct behavioral deficits relevant to motor learning and cued fear conditioning.

The first weeks of postnatal striatal development is an important period of excitatory synaptogenesis onto SPNs ^27,28,54^ and the cellular composition of the striatum during this time has been understudied. We surprisingly found that neurogenic progenitors make up a large component of the early postnatal striatum and that deletion of *Foxp1* decreases the ratio of these neurogenic progenitors to mature SPNs. These findings suggest that Foxp1 regulates intermediate progenitor pools and the differentiation of SPNs within the developing striatum. Furthermore, we found that Foxp1 is required for the specification of iSPNs that localize to the matrix and striosome compartments. iSPNs that remain with deletion of *Foxp1* localize to the striosome-matrix border and significantly upregulate top marker genes of a recently identified eSPN population, including *Foxp2*^13^. Future work will help resolve the functional contribution of eSPNs to striatal development.

Deletion of *Foxp1* specifically within iSPNs leads to both cell-autonomous and non-cell-autonomous changes in SPN projection patterns. Fewer iSPN terminals onto the globus pallidus external and fewer dSPN terminals onto the globus pallidus internal were observed. dSPNs and iSPNs are known to form inhibitory axon collaterals onto neighboring SPNs and modulate their excitability^55,56,57^. iSPNs and dSPNs also cooperate together to intermix within the striosome and matrix compartments^58^. We not only found that manipulation of iSPNs led to functional changes of dSPNs, but we captured a molecular snapshot of this inter-SPN communication, including differentially expressed ASD-risk genes involved in neuron projection such as *Cntn4, Cdh8*, and *Kirrel3.*

*FOXP1* is among a subset of genes repeatedly and significantly linked to ASD^59,60^. Thus, connecting our molecular findings to behavioral deficits is particularly relevant to a behaviorally diagnosed disorder that hinges upon two key behavioral phenotypes, impairments in language and social interactions and restrictive or repetitive behaviors. The majority of individuals with *FOXP1* mutations are diagnosed with ASD and all reported cases are comorbid with intellectual disability, gross motor delays, and/or selective language impairments^30,31^. We found that mice with iSPN-deletion of *Foxp1* caused significant motor disruptions, as measured by increased hyperactivity and motor-learning deficits on the rotarod. Concordant with our data, mice with ablated iSPNs or mice with *Darpp32* deletion from iSPNs were also hyperactive in the open field^61,62^. Adult mice with induced ablation of D2-receptors displayed severe motor learning impairments on the accelerating rotarod^63^. These data indicate that loss of iSPNs with deletion of *Foxp1* lead to significant motor-learning and activity deficits.

Pup USVs are an important measure of affective state and social behavior in mice^51,64^ and peak between postnatal days 4 and 10^35^. Disruption of neonatal call number and structure with deletion of *Foxp1* within dSPNs is particularly interesting given the high co-expression of both Foxp1 and Foxp2 within this cell-type and the ability of Foxp1 and Foxp2 to heterodimerize to regulate gene expression^65^. Foxp2 plays a critical role in the vocal behavior across many species, including humans, mice and songbirds^66^. We show that *Cntnap2*, a known shared target of Foxp1 and Foxp2^47,48^, is significantly upregulated within dSPNs. Variants in *CNTNAP2* are also associated with ASD and *Cntnap2* KO mice have altered pup USVs^48^. We previously found that *Foxp1* heterozygous mice display altered USV phenotypes, including deficits in call number, call structure, and pitch^35^. Additionally, mice with cortical and hippocampal deletion of *Foxp1* also produced fewer USVs, though no changes were observed in call structure or pitch^39^. Here, we observed changes in all three parameters within *Foxp1*^*D1*^ and *Foxp1*^*DD*^ mice suggesting that Foxp1 regulates distinct aspects of mouse vocal behavior largely through cortical-striatonigral circuitry.

Striosome compartments are smaller and architecturally disorganized with deletion of *Foxp1* in iSPNs and/or dSPNs in the early postnatal striatum. Loss of striosome-matrix compartmentalization is particularly striking in adulthood with dSPN-specific deletion of *Foxp1*. These findings indicate that dSPN-targets regulated by Foxp1 exert a stronger influence over maintaining striatal neurochemical organization. Behaviors specific to *Foxp1*^*D*1^ mice include deficits in contextual fear memory recall, a known limbic-circuitry associated behavior. Striosomes receive preferential inputs from limbic subcortical regions, including the amygdala and bed nucleus of the stria terminalis^8^; thus, inputs from these limbic regions targeting striosomes may be disrupted and contribute to the limbic-associated behavioral deficits seen in *Foxp1*^*D1*^ and *Foxp1*^*DD*^ mice. Additionally, mice with cortical and hippocampal deletion of *Foxp1* did not show deficits in cued or contextual fear conditioning^38^. Therefore, Foxp1 is likely mediating fear conditioned behaviors via disruption of striatal circuits.

While ASD is a genetically complex disorder, several studies have shown that striatal SPNs may be particularly vulnerable to ASD-linked mutations^67–71^. Our study uncovers the molecular targets of Foxp1 in SPN subtypes and finds that Foxp1 regulates ASD-relevant behaviors via distinct striatal circuits. We show that iSPNs are particularly vulnerable with loss of Foxp1 and that Foxp1 regulated iSPN-targets are enriched for high-confidence ASD risk-genes, suggesting that striatopallidal circuitry might be particularly at risk with loss-of-function *FOXP1* mutations. Our data provide important molecular insights for the development of future therapies targeting striatal circuits.

## Acknowledgments

Our sincerest thanks to Dr. Helen Lai, Dr. Jane Johnson, Dr. Said Kourrich, Marissa Co, and Dr. Fatma Ayhan for providing critical feedback on the manuscript. G.K. is a Jon Heighten Scholar in Autism Research at UT Southwestern. This work was supported by NIH/NIMH (T32-MH076690) to A.A, the Simons Foundation (SFARI 573689 and 401220), the James S. McDonnell Foundation 21^st^ Century Science Initiative in Understanding Human Cognition – Scholar Award (220020467), the Chan Zuckerberg Initiative, an advised fund of Silicon Valley Community Foundation (HCA-A-1704-01747), and grants from the NIH (DC014702, DC016340, MH102603) to G. K. We thank Dr. Denise Ramirez, Dr. Julian Meeks, and Apoorva Ajay from the UT Southwestern Whole Brain Microscopy Facility (WBMF) for assistance with volumetric whole brain imaging and automated image analysis. The WBMF is supported by the Texas Institute of Brain Injury and Repair and the UTSW Peter O’Donnell, Jr. Brain Institute. We thank the Neuroscience Microscopy Facility, supported by the UT Southwestern Neuroscience Department and the UTSW Peter O’Donnell, Jr. Brain Institute. We would also like to thank Dr. Shari Birnbaum at the UTSW Rodent Behavior Core for help performing the fear conditioning, novel-cage activity, and digigait analyses.

## Author contributions

A.A and G.K designed the study and wrote the paper. A.A performed all single-cell sequencing experiments and library preparations. A.A performed all RNA/protein quantification analyses, immunohistochemistry experiments, and mouse behavior. A.K developed the manual pipeline for scRNA-seq analysis. A.A and A.K analyzed the data. M.H contributed to mouse husbandry and genotyping.

## Methods

### Mice

All experiments were performed according to procedures approved by the UT Southwestern Institutional Animal Care and Use Committee. *Foxp1*^*flox/flox*^ mice^72^ were provided by Dr. Haley Tucker and backcrossed to C57BL/6J for at least 10 generations to obtain congenic animals as previously described^38,39^. *Drd1a-Cre* (262Gsat, 030989-UCD) and *Drd2-Cre* (ER44Gsat, 032108-UCD) mice were obtained from MMRC. *Drd2-eGFP* ^36^ and *Drd1-tdTomato*^73^ mice were provided by Dr. Craig Powell. We bred individual *Cre* or reporter lines to *Foxp1*^*flox/flox*^ mice to obtain all *Foxp1 cKO* mice in one litter that were heterozygous for *Cre* or reporter transgene. Mice used for single-cell RNA-sequencing and behavior experiments were not crossed with *Drd1-* or *Drd2*-reporter mice. Reporter mice were crossed with *Foxp1 cKO* lines for immunohistochemistry experiments and neuronal projection quantification. Mice were maintained on a 12-hr light on/off schedule.

### Protein isolation and immunoblotting

Striatal tissue was dissected, flash frozen, and stored at −80C before protein extraction. Protein was extracted from tissue using 1X RIPA Buffer (750mM NaCl, 250mM Tris-HCl pH7.4, 0.5% SDS, 5% Igepal, 2.5% Sodium deoxycholate, 5mM EDTA, 5mM NaVO4) with fresh protease inhibitor cocktail (10ul/ml), 10ul/ml of 100mM PMSF, and 25ul/ml of 200mM sodium orthovanadate. Tissue was homogenized in RIPA buffer using the TissueLyser LT (Qiagen) with a sterile, stainless-steel bead for 1min at 50 Hz. Samples were agitated for 1hr at 4C, spun down at 12,000rpm for 15 min, and supernatant was transfer to a fresh tube. Protein was quantified using a standard Bradford assay (Bio-Rad) and 20ug of protein per sample were run on 10% SDS-Page gels. PVDF membranes (Bio-Rad, 162-0177) were incubated in blocking solution (1% Skim milk in TBS with 0.1% Tween-20) for 30 min at room temperature (RT) and probed with primary antibodies overnight at 4C. Membranes were washed with TBS-T (TBS with 0.1% Tween-20) and incubated with appropriate, species-specific fluorescent secondary antibodies in blocking solution for 1hr at RT, and washed in TBS-T. Images were collected using the Odyssey infrared imaging system (LI-COR Biosciences).

### RNA isolation and quantitative real-time PCR

RNA from fresh or flash frozen tissue was harvested using miRNAeasy kit guidelines. RNA was converted to cDNA using recommended guidelines from SSIII Superscript Kit (Invitrogen) and qRT-PCR was performed using the CFX384 Real-Time System (Bio-Rad).

### Immunohistochemistry

For P7 or P9 mice, rapid decapitation was performed. Brains were extracted and dropped into ice-cold PBS for 1min before transfer into 4% PFA overnight. Brains were then transferred to 30% sucrose for 48 hours. 35um coronal slices were made using a SM2000 R sliding microtome (Leica) and free-floating sections were stored in PBS with 0.01% sodium azide. Slices were washed with TBS and incubated for 30min in 3% hydrogen peroxide in PBS, washed, then incubated in 30min in 3M glycine in 0.4% Triton-X, TBS. Slices were incubated in primary antibodies overnight at 4C, washed, and incubated in secondary antibodies for 1hr at room temperature. Slices were washed then mounted onto slides and allowed to dry overnight. Sections were incubated in DAPI solution (600nM in PBS) on the slide for 5 minutes and washed 3X with PBS. Sections were allowed to dry before mounting coverslips using Prolong Diamond Antifade Mountant.

### Imaging and Analysis

Images were collected using a Zeiss Confocal laser scanning microscope (LSM880) and all image quantification was performed using Fiji image processing package. For iSPN quantification, 20X z-stack images of dorsolateral, dorsomedial, and ventral striatum were taken within one hemisphere of four separate striatal sections from anterior to posterior per animal (3 images/section, 4 sections/animal, at least 3 animals/genotype). All images were taken within approximately similar sections across samples. Maximum projection images were quantified within a 1024×1024 pixel field of view across all images and averaged per section. For striosome quantification, 10X z-stack images were taken from one hemisphere of four separate striatal section from anterior to posterior per animal (4 sections/animals, at least 3 animals/genotype). Individual MOR+ patches were numbered, and area measurements summed for the total striosomal area measurement per section. Total striatal area was also measured per section to calculate the percentage of striosome area to total area per section. Differences between genotypes were assessed using a one-way ANOVA with multiple comparisons.

### Antibodies

The following primary antibodies were used for either immunoblots (IB) or immunohistochemistry (IHC) experiments: chicken anti-GFP (1:1,000, Aves Labs, GFP-1010), rabbit polyclonal anti-MOR (1:350, Millipore, AB5511), rabbit polyclonal anti-PDE1A (1:500, Proteintech, 12442-2-AP), rabbit polyclonal anti-DARPP32 (1:1,000, Millipore, AB1778), goat anti-tdTomato (1:500, LifeSpan Biosciences, LS-C340696), mouse monoclonal anti-FOXP1 (1:500, Abcam, ab32010), rabbit polyclonal anti-FOXP1 (IHC:1:1,000, IB: 1:5,000^74^), rabbit polyclonal anti-Calbindin (1:500, Millipore AB1778), goat anti-FOXP2 (N-terminal) (1:500, Santa Cruz 21069), rabbit polyclonal anti-β-Tubulin (IB: 1:10,000, Abcam, ab243041), and mouse monoclonal anti-SOX4 (1:500, Abcam, ab243041). All IHC following secondary antibodies were used at a 1:1,000 dilutions Alexa Fluor 488 Donkey Anti-Chicken IgG (Thermo Fisher, 703-545-155), Alexa Fluor 555 Donkey Anti-Goat IgG (Thermo Fisher, A-21432), Alexa Fluor 647 Donkey Anti-Rabbit IgG (Thermo Fisher, 711-605-152), Alexa Fluor 647 Donkey Anti-Mouse IgG (Thermo Fisher, A-31571). For IB, the following secondary antibodies were used at a 1:10,000 dilution: IRDye 800CW Donkey anti-Rabbit IgG (Licor, 925-32213) and IRDye 680RD Donkey anti-Rabbit IgG (Licor, 925-68071).

### Tissue processing for single-cell RNA-sequencing (scRNA-seq)

Mice (P9) were sacrificed by rapid decapitation and brains were quickly removed and placed in ACSF (126mM NaCl, 20mM NaHCO_3_, 20mM D-Glucose, 3mM KCl, 1.25mM NaH_2_PO_4_, 2mM of CaCl_2_ and MgCL_2_ freshly added) bubbled with 95%O_2_ and 5%CO_2_. Coronal slices at 500um were made using a VF-200 Compresstome in ACSF and transferred to a recovery chamber at room temperature in ACSF with 50uM AP5, 20uM DNQX, and 100nM TTX (ACSF+cb)^75^. Striatal punches were taken from these slices and incubated in 1mg/ml of pronase in ACSF+cb for 5min. Punches were washed with ACSF+ 0.04% BSA twice and gently dissociated into single-cell suspension using polished Pasteur pipettes with 600um, 300um, and 150um opening diameters, sequentially. Cells were centrifuged and washed twice, filtered through Flowmi Tip 40uM strainers, and resuspended with ACSF+ 0.04% BSA. Cell viability was quantified using the trypan blue exclusion method and cell concentration was adjusted for targeted sequencing of 10,000 cells/sample using the 10X Genomics Single Cell 3’ Reagent Kits v2 protocol to prepare libraries^40^. A total of 16 mice (4 mice/genotype, 2 males and 2 females per genotype) were processed for single-cell sequencing. Libraries were sequenced using the McDermott Sequencing Core at UT Southwestern.

### Pre-processing of Sequencing Data

Raw sequencing data was acquired from the McDermott Sequencing Core at UT Southwestern in the form of binary base call (BCL) files. BCL files were then de-multiplexed with the 10X Genomics i7 index (used during library preparation) using Illumina’s bcl2fastq v2.17.1.14^76^ and *mkfastq* command from 10X Genomics CellRanger v2.1.1 tools^40^. Extracted paired-end fastq files (26 bp long R1 - cell barcode and UMI sequence information, 124 bp long R2 - transcript sequence information) were checked for read quality using FASTQC v0.11.5^76^. R1 reads were then used to estimate and identify real cells using *whitelist* command from UMI-tools v0.5.4^77^ program. A whitelist of cell-barcodes (putative real cells) and R2 fastq files were later used to extract reads corresponding to real cells only (excluding sequence information representing empty beads, doublets, low quality/degrading cells, etc.) using *extract* command from UMI-tools v0.5.4^77^. This step also appends the cell-barcode and UMI sequence information from R1 to read names in R2 fastq file. Extracted R2 reads were then aligned to reference mouse genome (MM10/GRCm38p6) from UCSC genome browser^78^ and reference mouse annotation (Gencode vM17) using STAR aligner v2.5.2b^79^ allowing up to 5 mismatches. Uniquely mapped reads were then assigned to exons using *featureCounts* program from Subread package (v1.6.2)^80^. Assigned reads sorted and indexed using Samtools v1.6^81^ were then used to generate raw expression UMI count tables using *count* command from UMI-tools v0.5.4^77,82^ program. This raw expression matrix contains cells as rows and genes as columns and can be further used for downstream analysis such as normalization, clustering, differentially expressed genes, etc.

### Clustering Analysis

Raw single-cell RNA-seq UMI count data was used for clustering analysis using Seurat R analysis pipeline^83^. First, cells with more than 50,000 molecules (nUMI per cell) and cells with more than 10% mitochondrial content were filtered out to discard potential doublets and degrading cells. Also, genes from mitochondrial chromosome and chromosomes X and Y were removed as samples were from mixed genders. This dataset is referred to as *primary filtered dataset*. Post filtering, the raw UMI counts from primary filtered dataset were used for log-normalization and scaled using a factor of 10,000 and regressed to covariates such as number of UMI per cells and percent mitochondrial content per cell as described in Seurat analysis pipeline^83^. To further identify the top variable genes, the data were used to calculate principal components (PCs). Using Jackstraw analysis, statistically significant PCs were used to identify clusters within the data using original Louvain algorithm as described in Seurat analysis pipeline followed by visualizing the clusters with uniform manifold approximation and projection (UMAP) in two dimensions^84^. Genes enriched in each cluster compared to the remainder of the cells (adj. p-value <= 0.05 and log fold change >= 0.3) were identified as described in Seurat analysis pipeline. Genes corresponding to each cluster were used to identify the cell-type by correlating to genes expressed in previously published adult mouse striatal single cell data^13^. Cell-types were assigned to clusters based on (i) statistically significant enrichment of gene sets using the hypergeometric test (with a background of 7,500 genes, the number of expressed genes within our dataset) and (ii) expression weighted cell-type enrichment (EWCE) analysis ^41^ (https://github.com/NathanSkene/EWCE). Clusters that overlapped significantly with multiple cell-types were called for the most significant overlap (smallest Adj. P-value) and analyzed for expression of top marker genes of known cell-types. Cells from clusters that fell into neuronal categories (referred to as *secondary neuronal dataset*) were used to re-cluster the cells to define specific spiny projection neuronal sub-types using a similar approach as described above. Note that two small clusters (Clusters-21, 22) that corresponded to excitatory cortical neurons and a cluster with less than 30 cells total (Cluster-24) were excluded from the secondary neuronal dataset UMAP plots to focus on striatal cell-types.

### Differential Gene Expression (DEG) Analyses

#### Pairwise DEG analysis SPNs

For the spiny projection neuronal sub-type clusters identified using secondary neuronal dataset, pairwise differential gene expression analysis tests were performed within each cluster-pair using a Poisson likelihood ratio test from the Seurat R analysis pipeline^83^ to identify genes enriched (adj. p-value <= 0.05, log_2_FC>|0.25|) in SPN sub-types.

#### Pseudobulk DEG analysis

Within the secondary neuronal dataset, neurons identified as either dSPNs (*Drd1*+) or iSPNs (*Drd2*+) were combined into pools of cells segregated by genotypes. Differential expression within pools of dSPN or iSPNs of Foxp1 cKO samples were then compared to control samples using Poisson likelihood ratio test from the Seurat R analysis pipeline accounting for averaged expression differences in either dSPNs or iSPNs across genotypes irrespective of the identified clusters. Significant expression changes (adj. p-value <=0.05, log_2_FC>|0.3|) reflected the differences in expression of genes in one specific cell population (dSPNs or iSPNs) across genotypes instead of detected clusters.

### Down-sampled Dataset Analysis

Cells from the primary filtered dataset were used to randomly select the cells from each genotype matching the number of cells present in each genotype with the lowest representation of the cells (*Foxp1*^*CTL*^ = 14466 cells, *Foxp1*^*D1*^= 16,961 cells, *Foxp1*^*D2*^ = 9,898 cells, *Foxp1*^*DD*^= 21,453 cells, using random sampling, the same number of cells from *Foxp1*^*CTL*^, *Foxp1*^*D1*^and *Foxp1*^*DD*^ were matched to *Foxp1*^*D2*^). This is referred to as the *primary down-sampled dataset*. This dataset was further used to separate the cells into clusters and identify cell-types as described in the clustering analysis section above. Clusters corresponding to SPNs from the primary down-sampled dataset (referred to as the *secondary down-sampled neuronal dataset*) were re-clustered to identify SPN subtypes in a similar manner as described in the clustering section above.

### Availability of Data and Code

The sequencing data reported in this paper can be access at NCBI GEO with accession number GSE125290. Code that was used to perform data pre-processing, clustering and differential gene expression analysis is available at GitHub repository (https://github.com/konopkalab/early-postnatal-striatal-single-cell-rna-seq).

### TissueCyte Imaging and Quantification

#### STPT and image acquisition

Serial two-photon tomography (STPT)^49^, in which automated block face imaging of the brain is repetitively alternated with vibratome sectioning, was conducted on the TissueCyte 1000 platform using the manufacturer’s custom software for operation (Orchestrator). Mouse brains were perfusion-fixed in 4% paraformaldehyde and embedded in low-melting point oxidized agarose (4.5% w/v; Sigma #A0169). Vibratome sections were prepared at 75 µm thickness using a frequency of 70 Hz and a speed of 0.5 mm/sec. 185-190 total sections were collected of each brain. A 9 by 13 mosaic of tile images was collected at each level using lateral resolution of 0.875 µm/pixel. Optical sectioning was used to collect three z-planes within each 75 µm physical section to obtain 25 µm axial resolution. The two-photon excitation laser (Spectra Physics MaiTai DeepSee) was tuned to 920 nm to excite both eGFP and tdTomato. The emission fluorescence from the red, green and blue channels was independently collected using photomultiplier tube detectors. The tile images were saved to network attached servers and automatically processed to perform flat field correction and then stitched into single-channel 2D coronal sections in 16-bit .tif format using the manufacturer’s custom software (AutoStitcher).

#### Sample preparation and details

Mice (8 weeks) were perfused with PBS followed by 4% PFA. Brains were removed and post-fixed overnight in 4% PFA at 4C. Samples were transferred to PBS + 0.1% sodium azide and stored at 4C until imaging. A total of 19 whole mouse brain images were collected in three cohorts for machine learning analysis according to their patterns of fluorophore expression. The first cohort consisted of 8 samples expressing tdTomato (detected predominantly in the red channel), the second cohort had 8 samples that expressed eGFP (detected predominantly in the green channel) and the third cohort consisted of 3 dual-labeled (eGFP + tdTomato) samples.

#### TissueCyte image processing and registration

STPT image processing was performed via BioHPC, an advanced computing cluster at UT Southwestern. All channels of the coronal sections were downsampled to 10 µm lateral resolution, intensity adjusted to fill the 16-bit range, and combined to form 3D image stacks using custom MATLAB software. The image stacks were then processed through a 3D median filter to remove high-contrast noise. The 3D image stacks were registered to Allen Institute for Brain Science Common Coordinate Framework (version 3, CCFv3) at 10 µm × 10 µm × 100 µm resolution using NiftyReg software^85^. Briefly, registration involved three steps: (i) Affine transformation (reg-aladin) for global registration (ii) Cubic B-spline transformation (reg-f3d) to achieve local transformation and (iii) Resampling the transformed brains to Atlas coordinates (reg-resample). Registration transformations were established based on the red channel, then applied equally to all other data channels, including the probability maps (described below).

#### Interactive Image training for classifying signals of interest

The three raw channels of the 2D stitched coronal sections were downsampled to 1.5 µm lateral resolution. A maximum intensity projection of the three optical sections was produced for each physical section across all 3 color channels, creating an RGB image stack with the same number of 2D frames as physical sections (e.g. 185 or 190). Ilastik (Interactive learning and segmentation toolkit)^50^ software was deployed on BioHPC and used to train a pixel-wise random forest classifier to identify features of interest (e.g. fluorescent neuronal cell bodies and axonal projections). Three or four representative sections were chosen from the 185-190 image stack for model training. A supervised random forest model was trained by users to classify fluorescent features of interest (e.g. eGFP and/or tdTomato), and to distinguish them from other image features (e.g., bright microbubbles, empty space, autofluorescence) using the interactive features in Ilastik. An independent random forest model was trained for each of the image batches described above. The random forest classifiers were used to detect features of interest in all image sections, creating a “probability map” for each voxel in each 3D whole brain image. In these probability map images, the value of each voxel in each virtual channel (corresponding to each image feature, e.g. eGFP) represents the probability that the voxel includes information for the desired feature. These exported probability maps were registered to the CCFv3.0 using the transformation parameters using NiftyReg (reg-aladin).

#### Quantification and visualization

The features of interest in the registered probability maps were quantified by automatically segmenting brain regions of interest based upon CCFv3.0 volumetric annotations. Custom MATLAB software aggregated brain regions of interest (e.i., nucleus accumbens, caudate putamen, globus pallidus external and internal, substantia nigra pars compacta and pars reticulata), calculated the cumulative probabilities of all voxels in each region, and normalized these values by the volume of each structure. This exported data matrix thus included normalized probability intensity values for each machine learning feature, each brain region of interest, and each brain. For visualization, the combined probability map stacks were rendered in 3D using the ClearVolume plugin for Fiji/ImageJ^86^.

### Behavior tests

#### Open Field

Mice age 8-12 weeks were allowed to acclimate to the testing room for 1hr before being placed in a 55cm x55cm × 36cm matrix (Phenome Technologies) and recorded for 30min. Total distance and velocity measurements were analyzed using Actimetrics LimeLight software.

#### Novel-cage activity

As previously described^38^, mice were moved into individual cages (18×28cm) with minimal bedding. Cage was placed into a dark Plexiglas box and the movements were measured using a Photobeam Activity System-Home Cage software for two hours. The number of beam breaks was recorded every 5 min and averaged over two hours for statistical analyses.

#### Rotarod

Following previously published methods^35^, mice (8-12 weeks) were acclimated to the testing room for 30min before placed in one lane of a 5-lane accelerating rotarod (Series 8 ITCC Life Science rotarod). The textured drum within the individual lanes was programed to accelerate from acceleration from 4-40 rpm within a maximum time frame of 300 sec. Each mouse was positioned facing away from the experimenter. Latency to fall was recorded once the trial was initiated. Manual activation of the sensors occurred when an animal made a full rotation holding onto the drum. Animals received four trials per day (20min intervals) with lanes cleaned between animals with NPD over the course of three consecutive days.

#### Grip strength test

Grip strength was tested following previously published methods^35^. Briefly, following rotorad experiments, the forelimb and hindlimb grip strength mice were measured using Chatillon Force Measurement equipment. The forelimbs, followed by the hindlimbs, for each animal were tested first by placing forelimb paws on a mesh wire meter and pulling them away from the wire at constant force. Five consecutive measurements were recorded for both hindlimbs and forelimbs and averaged for a final grip strength measurement.

#### Nestlet behavior

Nesting behavior was analyzed using a previously published approach^38,52^. Mice (8-12 weeks) were isolated into clean cages overnight with 3 g of intact nestlet. After 16-18 hrs, the amount of unused nestlet was measured and images of the nests were taken to assess the quality and given a score.

#### Neonatal ultrasonic vocalization measurements

USVs were recorded as described previously^35,38^. Briefly, pups were isolated from dams at P4, P7, and P10 and placed into a soundproof container. USVs were recorded for 3min with an UltraSoundGate condenser microphone using Avisoft Bioacoustic software. Analysis of sound spectrograms was automatically performed using MATLAB codes^87^.

#### Digigait

Mice (8-12 weeks) were placed onto the transparent treadmill using the DigiGait Imaging System (Mouse Specifics, Inc) at 10 cm/sec. The speed was quickly increased to 20 cm/sec with a high-speed video camera mounted under the clear treadmill to capture images of all four paws at the 20 cm/sec speed. A section of video with at least 6-10 steps is analyzed and the paw placement is automatically detected and quantified by the software system. Right and left forelimb and hindlimb paw measurements were analyzed separately.

#### Fear Conditioning

Fear conditioning was measured using boxes with metal grid floors connected to a scrambled shock generator (Med Associates Inc., St. Albans). Mice were trained by placing them individually in the chamber for 2min before they received 3 tone-shock pairings (30sec white noise, 80dB tone, co-terminated with a 2 sec, 0.5mA footshock, 1min intertrial interval). Twenty-four hours later, contextual memory was measured by placing the mice into the same chamber and measuring freezing behavior using the Med Associates software. Forty-eight hours post training, memory of the white noise cue was measured by placing mice in new environment, with altered floors, walls, different lighting, and a vanilla smell. Freezing was measured for 3 min and then noise cue was turned on for an additional 3 min and freezing was measured.

### Statistics and reproducibility

Statistical methods and code used for scRNA-seq and analysis are provided in the above methods sections. All statistical test used (and p-values obtained) for SPN projection analysis, behavior, and immunohistochemistry are described in figure legends. No statistical methods were used to estimate sample size, but behavior cohorts were based on previously published papers ^35,38,39^. Sample size for each experiment is indicated in figure legends.

---

**Supplementary Figure 1.**
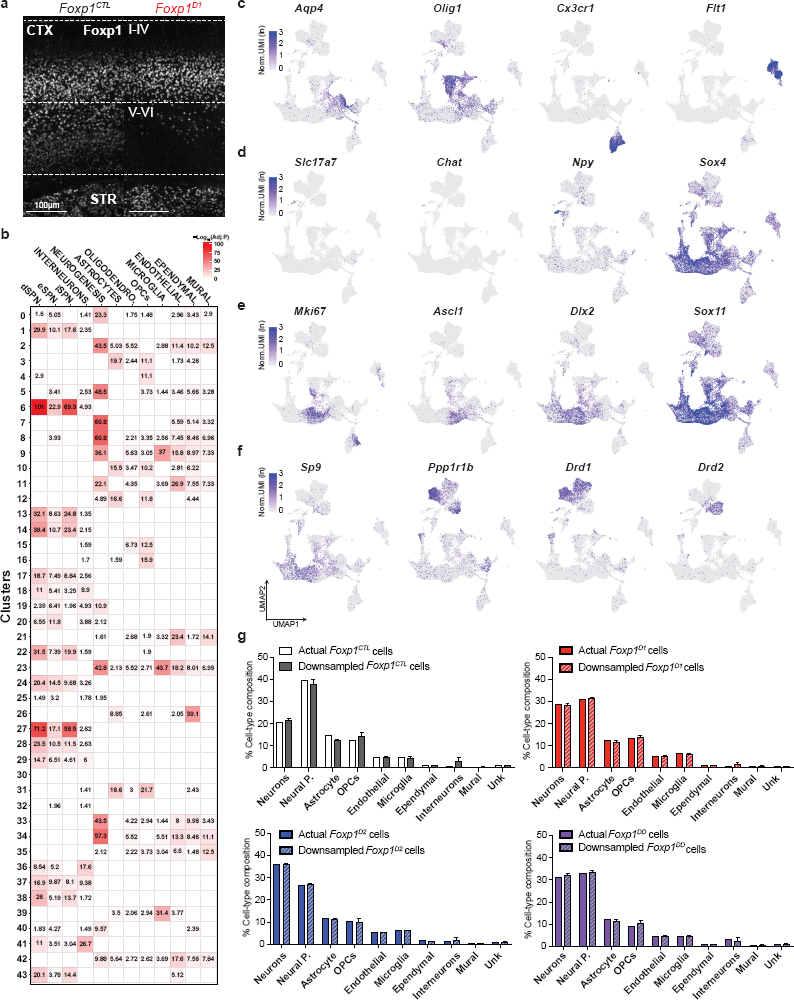
Cell-type annotation of early postnatal striatal scRNA-seq. **a**) Confocal imaging of the somatosensory cortex of *Foxp1*^*CTL*^ or *Foxp1*^*D1*^ adult mice showing reduction of Foxp1 expression within cortical layers V-VI (scale bar is 100um, CTX= cortex, STR=striatum). **b**) Heatmap showing the enrichment of genes within each cluster that correlate to a previously annotated dataset (Saunders et al., 2018) using the hypergeometric overlap test. **c**-**d**) Expression plots showing the normalized UMI (ln) for known marker genes of distinct cell-types: (**c**) *Aqp4* for astrocytes, *Olig1* for OPCs, *Cx3cr1* for microglia, *Flt2* for endothelial, (**d**) *Slc17a7* for glutamatergic cortical neurons, interneuron populations (*Chat, Npy*), neurogenic and neural differentiation marker (Sox4). (**e**) Expression plots of markers identifying neurogenic populations: proliferating cells (*Mki67*), neural progenitors (*Ascl1*), neural progenitors derived from the lateral ganglionic eminence (*Dlx2*), neurogenic and neural differentiation marker (*Sox11*). **f**) Expression plots of genes important for iSPN specification (*Sp9*), mature SPN marker (*Ppp1r1b*), and major SPN subtypes (*Drd1, Drd2*). **g**) No changes in cell-type composition were observed between the average down-sampled datasets (10 iterations with 9,898 cells within each genotype) compared to the actual dataset.

**Supplementary Figure 2.**
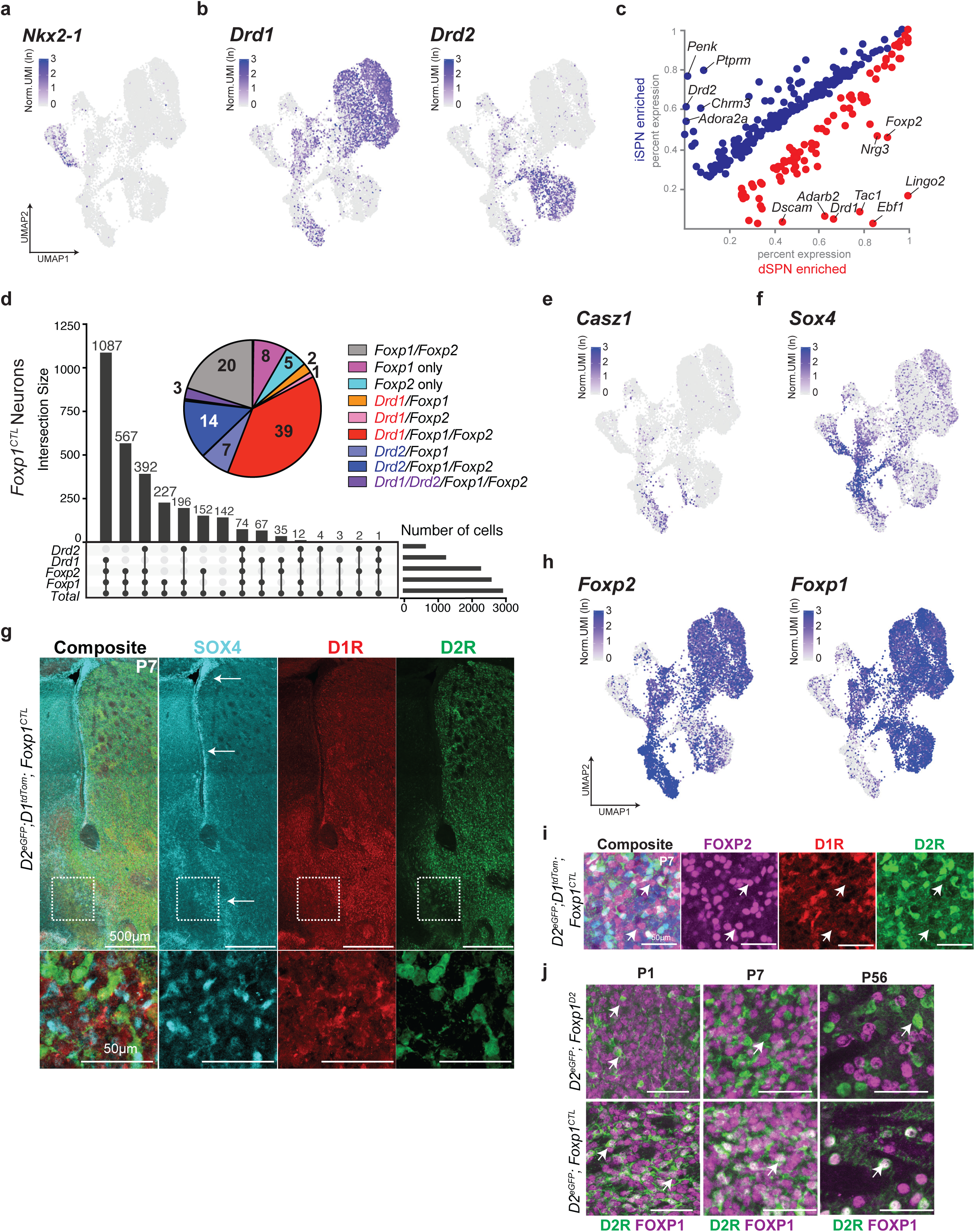
Neuronal subclusters and the intersection of Foxp1 and Foxp2 expressing striatal neurons. **a**-**b**) Expression plots with the normalized UMI counts for interneuron marker *Nkx2-1* (**a**) or SPN markers *Drd1* (dSPNs) or *Drd2* (iSPNs) (**b**). **c**) Scatter plots showing the percent expression of enriched transcripts between the largest iSPN (Cluster-2) and dSPN (Cluster-0) clusters. **d**) Upset plot showing the number of cells that overlap in expression of *Drd1, Drd2, Foxp1*, or *Foxp2* transcripts within neurons of control samples. Pie chart inlet shows the percent composition of this overlap (percentages <1% not visualized). **e**-**f**) Expression plots with the normalized UMI counts for eSPN marker *Casz1* (**e**) and imSPN marker *Sox4* (**f**). **g**) Coronal striatal image of control animals crossed to both *Drd1-tdTomato* and *Drd2-eGFP* reporter mice to label dSPNs or iSPNs, respectively, and stained for Sox4 at P7 (500μm scale bar). White arrows indicate the location of Sox4+ neurons, with inlet showing 63X confocal image (50μm scale bar). **h**) Expression plots with the normalized UMI counts for *Foxp2* and *Foxp1*. **i**) The same mice from (**g**) stained for Foxp2. White arrows indicate example cells where Foxp2 does not co-localized with either dSPNs or iSPNs (50μm scale bar). **j**) Foxp1 is not expressed within remaining iSPNs within *Foxp1*^*D2*^ mice crossed to Drd2-eGFP reporter mice at P1, P7, or P56 (adult) timepoints (50μm scale bar).

**Supplementary Figure 3.**
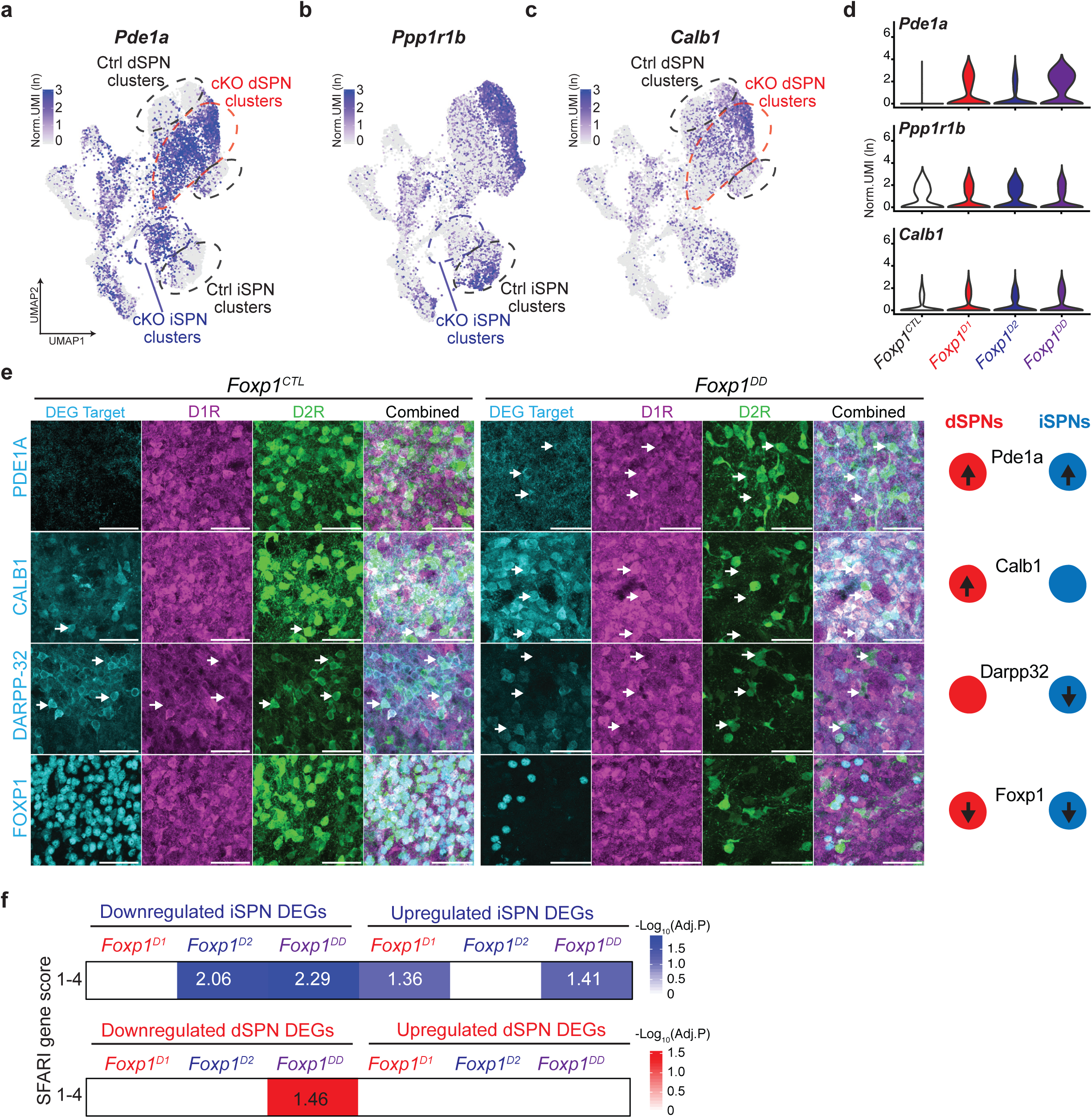
Confirmation of cell-type-specific targets regulated by Foxp1 and overlap with ASD-associated genes. **a**-**c**) Expression plots of significant DEGs regulated by Foxp1 in both iSPNs and dSPNs (*Pde1a*), iSPNs (*Ppp1r1b*), or dSPNs (*Calb1*). **d**) Violin plots showing the average normalized UMI (ln) of significant DEGs across genotype within all neuronal clusters of *Pde1a Ppp1r1b*, and *Calb1*. **e**) 63X confocal images of coronal, striatal sections stained for Pde1a, Calb1, and Darpp32 in *Foxp1*^*CTL*^ and *Foxp1*^*DD*^ mice crossed to reporter mice labelling dSPNs with tdTomato and iSPNs with eGFP (50μm scale bars). White arrows indicate specific cells where Foxp1 is either 1) upregulating a target (Pde1a) in both dSPNs and iSPNs, 2) upregulating a target (Calb1) in dSPNs only, or 3) downregulating a target (Darpp32) in iSPNs only. **f**) Enrichment of ASD-risk genes SFARI score 1-4 with upregulated or downregulated iSPN-DEGs (blue) or dSPN-DEGs (red) across *Foxp1 cKO* samples using a hypergeometric overlap test (8,000 genes used as background).

**Supplementary Figure 4.**
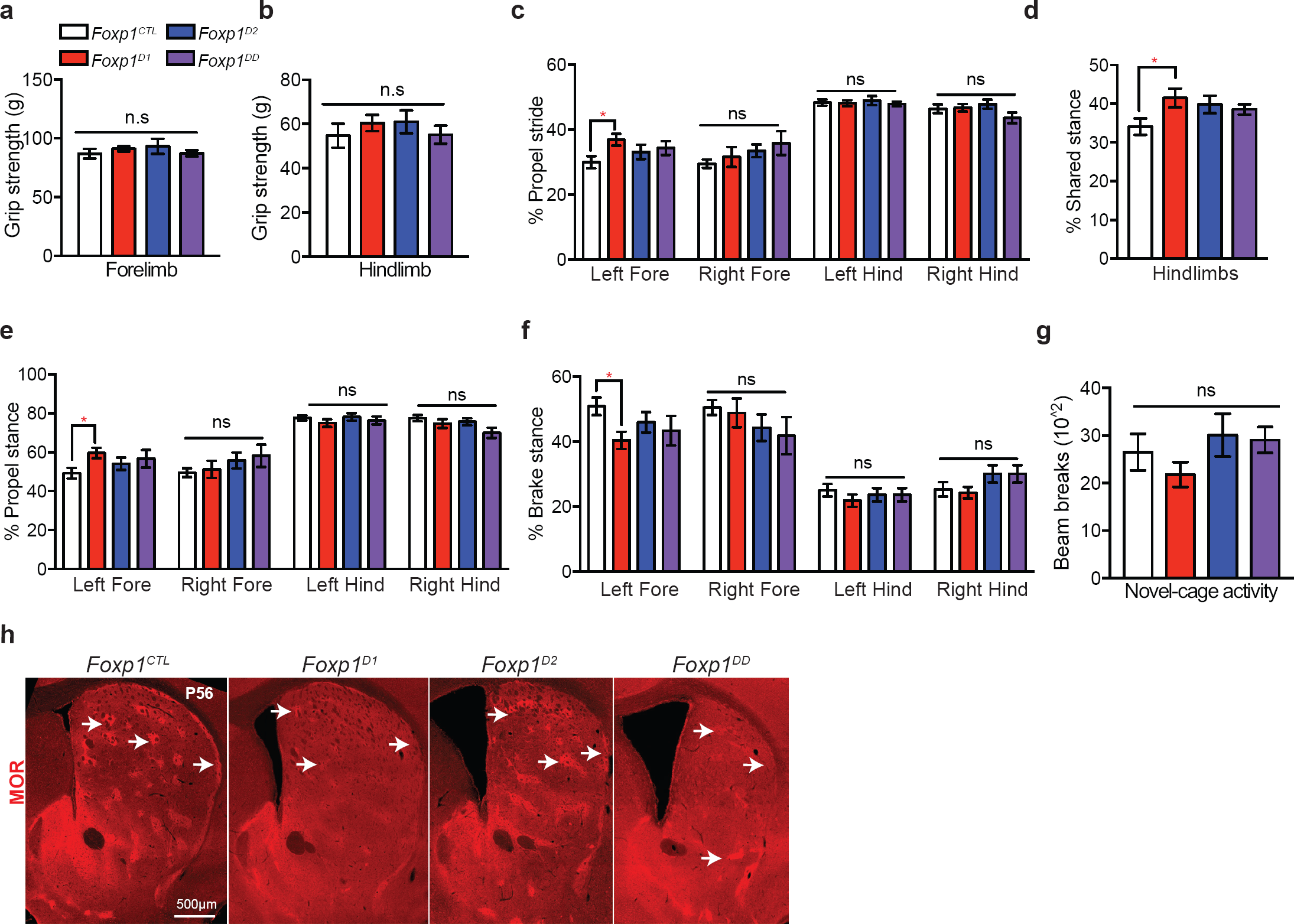
Supplemental behavioral testing of *Foxp1 cKO* mice. **a**-**b**) No change in forelimb (**a**) or hindlimb (**b**) grip strength was detected across *Foxp1 cKO* mice. Data are represented as mean ± SEM. n=12 *Foxp1*^*DD*^; n=17 *Foxp1*^*D1*^; n=16 *Foxp1*^*D2*^; n=11 *Foxp1*^*CTL*^. Forelimb: p=0.8520 (*Foxp1*^*D1*^), p=0.6477 (*Foxp1*^*D2*^), p=0.999 (*Foxp1*^*DD*^); Hindlimb: p=0.7225 (*Foxp1*^*D1*^), p=0.6786 (*Foxp1*^*D2*^), p=0.999 (*Foxp1*^*DD*^), one-way ANOVA with Dunnett’s multiple comparisons test. **c**-**f**) Digigait analysis examining propel stance (**c**), shared stance (**d**), propel stride (**e**), or brake stance (**f**) across *Foxp1 cKO* mice. Only *Foxp1*^*D1*^ mice exhibited a significant increase in left forelimb propel stance, propel stride, shared stance, and decrease in left forelimb break stance compared to control animals. Data are represented as mean ± SEM. n=7 *Foxp1*^*DD*^; n=9 *Foxp1*^*D1*^; n=10 *Foxp1*^*D2*^; n=10 *Foxp1*^*CTL*^. *p<0.05, two-way ANOVA with Dunnett’s multiple comparisons test. **g**) Activity levels within a novel-cage environment were unaltered in *Foxp1 cKO* mice. Data are represented as mean ± SEM. n=7 *Foxp1*^*DD*^; n=9 *Foxp1*^*D1*^; n=10 *Foxp1*^*D2*^; n=10 *Foxp1*^*CTL*^. p=0.6834 (*Foxp1*^*D1*^), p=0.8145 (*Foxp1*^*D2*^), p=0.9374 (*Foxp1*^*DD*^), one-way ANOVA with Dunnett’s multiple comparisons test. **h**) Confocal images of striatal sections stained for Mu-Opiod Receptor (MOR) across adult *Foxp1 cKO* mice. White arrows show example striosomes (500μm scale bar).

**Supplementary Table 1. Differentially expressed genes within clusters.** Significantly enriched genes within each cluster identified within the total scRNA-seq dataset (all-cells) and the neuron specific re-clustering dataset. Columns show the gene name, average log fold change, cluster affiliation, percent expressed within that cluster, adjusted p-value.

**Supplementary Table 2. Cluster cell-type affiliation and number of cells within each cluster across genotype**. The total number of clusters, total number of cells within each cluster across genotype, and cluster-cell-type affiliation are shown within respective columns for all-cells and neuron specific datasets.

**Supplementary Table 3. Pairwise differential gene expression analysis between dSPN and iSPN subclusters.** Differential gene expression analysis was performed between each main subcluster within control dSPNs (Clusters-0, 5, 9) or iSPNs (Clusters-2, 8).

**Supplementary Table 4. Pseudobulk differential gene expression analysis between individual Foxp1 cKO dSPNs or iSPNs relative to control dSPNs or iSPNs.** All significant, differentially expressed genes uncovered in the pseudobulk analysis within iSPNs or dSPNs across *Foxp1 cKO* samples relative to the same cell-type in control samples. D1cKOs= *Foxp1*^*D1*^ samples, D2cKOs= *Foxp1*^*D2*^ samples, DDcKOs= *Foxp1*^*DD*^ samples.

**Supplementary Table 5. Gene ontology analyses of DEGs within dSPNs or iSPNs found in *Foxp1 cKO* samples.** GO analysis of Foxp1 regulated DEGs using Toppgene (https://toppgene.cchmc.org).

